# Naturally arising *de novo* open reading frames as potential zinc chelators in *Drosophila melanogaster*

**DOI:** 10.64898/2026.08.19.745823

**Authors:** UnJin Lee

## Abstract

A central open problem in the study of *de novo* gene origination is that the molecular mechanisms and functions driving the emergence of such evolutionarily young, *de novo* protein-coding genes remain poorly understood. Metal chelation, a simple, directly selectable activity that both requires no specific interaction partners and is also compatible with intrinsic disorder, is one possible function. This possibility was tested using sequence signatures in 7,849 transcriptionally supported, still-segregating *Drosophila melanogaster de novo* open reading frames. Interestingly, these new open reading frames (neORFs) are enriched for bis-histidine motifs at the metal-coordination-competent spacings H-x-H and H-x-x-x-H and show no enrichment at the incompatible even spacings relative to repeat-masked intergenic ORFs. Notably, these neORFs were also found to lack the C-x-x-C grammar of canonical metal-binding proteins. I report that this H-x-H bis-histidine signal is generated by translation of (CA)ₙ microsatellites into His-Thr-His in *Drosophila melanogaster*, as evidenced by a CAC-codon bias within H-x-H motifs and a fourfold threonine enrichment at the central position. I propose that recurrent microsatellite expansion supplies *Drosophila* with a distributed, independently originated class of candidate metal-binding *de novo* peptides. I then trace one neORF (ZMEG) from conserved ancestral non-coding sequence to a transcribed, *melanogaster*-lineage open reading frame whose (CA)9-derived His run presents a candidate His32–His36 bis-histidine site. I propose that such *de novo* proteins may constitute a class of molecules united not by common descent but by their shared origin in evolvable repeat sequence, highlighting the importance of emergence bias in molecular evolution.

**Significance Statement:** Where do brand-new genes come from, and what do they do? Most genes are inherited and modified over eons, but some protein-coding genes arise de novo from previously non-coding DNA, and their functions have remained mysterious. Analyzing thousands of young Drosophila genes, I find that an abundant, highly mutable class of repetitive DNA, (CA)n microsatellites, has the potential to encode repeating histidine pairs capable of grabbing metal ions such as zinc. This links a simple, error-prone genomic feature to a concrete, selectable biochemical activity, and traces one such gene from non-coding ancestry to a candidate metal-binding peptide. The findings suggest biochemically active proteins can evolve not only by copying old genes but also by exploiting the predictable emergence of repeat sequences.

---

While the majority of work on new gene origination, or the appearance of evolutionarily young genes, has been focused on duplicate gene evolution and gene family expansions (*1*), much effort has been made to both establish and validate the existence of genes that have arisen *de novo*, i.e. protein-coding genes that have arisen from ancestrally inactive genomic sequences (*2*). Early work focused particularly on understanding transcription of such genes on both a species (*3*) and population (*4*) level within *Drosophila*, while later efforts utilized ribosomal profiling techniques to demonstrate widespread translation of *de novo* open reading frames (ORFs) (*5*, *6*). In parallel, a small number of well-characterized examples of *de novo* genes have been described across a number of clades, including the anti-freeze glycoprotein (AFGP) in codfish (*7*), Goddard in *Drosophila* (*8*, *9*), Pldi in mouse (*10*, *11*), NCYM in humans (*12*), and more. Despite lingering doubts, subsequent follow-up work has been performed in order to characterize the biophysical properties of such *de novo* genes (*13–15*), revealing common trends, such as high levels of intrinsic disorder and short ORF length (*15*) — effects that have generally been ascribed to both pre-adaptation (*14*, *16*), which is the idea that pervasive translation exposes variants to selection prior to emergence, and the ambush hypothesis (*17*), which is the idea that frame-shifted codons are positively selected to terminate off-frame translation. Despite this work, widespread acceptance of the existence of *de novo* genes has been hampered by the lack of ascribable function for these genes.

Prior efforts have attempted to explain the evolutionary forces driving the origination and retention of these genes, such as 1) the proto-gene model (*18*), where ORFs sit on a continuum between non-genic and fully genic states, 2) the transcription-first model (*2*), where transcription precedes gain of translation, and 3) the pre-adaptation model (*14*, *16*). However, such models are generally unsatisfying in that they do not directly tackle the question of function. One exception is the recently proposed cultivator model of *de novo* gene origination (*19*, *20*), which posits that the transcriptional activation of a new promoter is a crude but highly evolvable mechanism that selects for transcriptional activity not of the newly evolved *de novo* transcript, but of its neighboring genes through physical, distance-based effects such as transcriptional interference (*21–23*) or supercoiling mediated coupling (*24*). Indeed, such rapid evolvability has been shown to be critical in the understanding of the evolutionary trajectories of duplicate genes arising via some form of emergence bias (*25*), either through the innovation-amplification-divergence model (*26*, *27*), where weak enzymatic side-activity is amplified via dosage, allowing fixed copies to then diverge, and/or the enhancer capture-divergence model (*28*), where a new gene copy inherits expression by capturing a pre-existing regulatory contact followed by divergence. However, the cultivator model remains unsatisfying, as it can only provide an evolutionary mechanism describing the earliest stages of *de novo* gene origination, i.e. non-coding RNA (ncRNA), and remains unsatisfactory for explaining the persistence and function of protein-coding *de novo* genes.

The general acceptance of *de novo* origination has also been further hampered by key confounding technical factors found in many *de novo* gene studies, e.g. short peptide length (*15*), and the natural limits of homology identification in divergent species (*29*). Instead, putative *de novo* gene origination events are difficult to untangle from the duplication-based origination mechanisms underlying orphan genes (*30*) — duplicate genes that have diverged so rapidly from their parental gene that homology is impossible to detect (*29*). However, the recent explosion of high-quality genome annotations of a vast number of species has accelerated the study of *de novo* genes, providing increased support for claims of *de novo* origination beyond the absence of detectable homology. For example, combined phylostratigraphic and transcriptomic methodologies heavily rely on synteny to *positively* demonstrate the absence of transcriptional units in ancestral states (*13*, *18*, *31*). While such approaches do not necessarily exclude the possibility of duplication, either symmetrically in tandem or asymmetrically over larger distances that traverse across topologically associating domains (TADs), they do allow for the positive identification of an absence of a transcription unit *at an ancestral locus*. Other efforts have also utilized direct identification of ancestral sequence in combination with synteny and transcriptomics to positively identify the lack of transcription via the usage of mock annotations in outgroup species (*20*). By using RNA-seq data from outgroup species, such an approach can definitively demonstrate that sequences ancestral to *de novo* genes (used here to include both protein-coding and ncRNA genes) were truly transcriptionally inactive (*20*). However, this approach has been used mostly to identify *de novo* non-coding RNAs, leaving a gap in our understanding of how *de novo* protein coding genes may arise naturally.

## What do de novo proteins do?

A comprehensive analysis of protein function provides the following list of possible mechanisms driving *de novo* gene fixation. First is pre-adaptation, the claim that selection acts to minimize the genomic harm done by previously fixed *de novo* proteins subsequent to the fixation of a random protein-coding gene (*14*, *16*). Unlike the other putative functions listed here, this function — or lack thereof — is driven by negative selection against some unknown deleterious interaction. Another possibility is the appearance of a fortuitous but beneficial interaction with some number of pre-existing interaction partners that drives the new *de novo* protein to fixation. Possible sub-mechanisms include general protein-protein interaction via disordered regions (*32*), the refinement of regulatory function of pre-existing genes (*19*), or the titration of thermodynamic properties of liquid-liquid phase separated condensates (LLPS). This last possibility also opens the door to more interesting possibilities, allowing for the potential of *de novo* proteins playing roles in cellular-scale phenomena either via direct interaction with condensates or surface adsorption (*33*). One concrete example would be similar in nature to the duplicate gene HP6/Umbrea, a known heterochromatin protein (*34*, *35*). Another potential mechanism would be membrane-based interactions (*36*). Whatever the case, protein-protein interactions often have strong sequence-specific signatures in the form of short linear motifs (SLiM)(*37*), such as in the case of the known PxVxL signature found in HP6/Umbrea (*34*).

One previously under-appreciated possibility is that *de novo* proteins may act as chelating agents, a function that is highly consistent with commonly observed properties of *de novo* genes such as intrinsic disorder. This is an attractive possibility, as chelation provides a simple, direct, selectable function that does not specifically require interaction between a nascent *de novo* protein and the rest of the proteome. While some degree of interaction is likely to occur between nascent *de novo* proteins and other proteins, variants producing such proteins are expected to be retained so long as the net selective effect of such incidental interactions remains positive. Further refinement or purging of such deleterious interactions could be explained by pre-adaptation (*16*), but the mechanism driving the original fixation events remains to be explained under that model.

Two primary signatures are known to be associated with chelation. First, pairs of cysteine residues in the configuration C-C or C-x-x-C are widely recurring motifs in metal- and disulfide-coordinating proteins (*38*). In *Drosophila*, proteins bearing these motifs, such as the cysteine-rich metallothioneins, are transcriptionally induced by the MTF-1 (metal-responsive transcription factor 1) pathway (*39*). Additionally, pairs of histidine molecules are also known motifs for chelation due to histidine’s partially protonated imidazole residue (*40*). The ordering for these histidines is critical for establishing metal-binding function, manifesting as H-x-H and H-x-x-x-H motifs, where x is an uncharged residue (*41*). Importantly, H-H and H-x-x-H motifs are disfavored for metal-binding function, as the imidazole residues chelate only when both residues converge on the same ion (*40*, *42*, *43*). Because histidine is itself the substrate of METTL9-family protein histidine methyltransferases, whose methylation can attenuate metal binding, any (CA)n-derived histidine site is in principle subject to post-translational control (*44*, *45*). In order to test for bis-histidine residue chelation as a mechanism driving *de novo* protein origination and retention, I utilized an existing dataset of high-confidence, population-level, new expressed ORFs (neORFs) to test for enrichment of histidine chelation-associated amino acid signatures in a panel of 7,849 transcriptionally supported, young *de novo* proteins that are still segregating in *D. melanogaster* (Materials and Methods)(*13*). Importantly, neORFs consist of the category of *de novo* ORFs that appear in fewer than 7 strains out of a total of 7 independent, iso-female lines of *D. melanogaster* (*13*). I then identify and analyze potential biochemical function of a candidate chelating neORF, which I term *zmeygorynych* (ZMEG), after the mythological three-headed dragon described in Eastern Slavic folklore.

## Results

### neORFs are enriched for H-x-H bis-histidine coordinate chemistry

To test for potential enrichment of bis-histidine-driven chelation in *de novo* proteins, the relative number of various H-(x)n-H configurations was calculated for three classes of validated genes: transcriptionally active neORFs (N=7,849, median length *ℓ*_0.5_=64 aa), transcriptionally inactive, intergenic, repeat-masked ORFs (N=1,472,446, *ℓ*_0.5_=22 aa), and known metal-binding proteins from UniProt (KW-0479)(N=856, *ℓ*_0.5_=538 aa). Specifically, the number of motifs of each category was calculated, then compared to the number of motifs expected by chance alone (Materials and Methods). An odd spacing between histidines is crucial for metal chelation because it allows for histidine’s partially protonated imidazole residues to coordinately bind free Zn^2+^ ions. In addition to a cyclic pattern of enrichment for odd n, the number of detected H-x-H patterns in neORFs was significantly enriched over the expectation from a composition-matched null expectation (Fisher’s Exact p=1.4×10^-9^; Figure 1A). Overall, 9.7% of the 7,849 de novo ORFs carry at least one bis-histidine H-x-H motif (0.194 motifs per 100 residues), while the intergenic control set has a 3.9% incidence rate (0.157 motifs per 100 residues). Additionally, no significant difference was found in the inert H-x-x-H configuration (Fisher’s Exact p=0.88). The relative number of neORFs again increases in the H-x-x-x-H configuration, then continues to repeat in a cyclical manner, consistent with the geometry of bis-histidine chelation. Notably, the relative number of inter-histidine residues reached a maximum at n=3 (H-x-x-x-H) for metal binding proteins, consistent with the α-helix register. This reflects different motifs predominant in HExxH zincin proteases, including the gluzincins thermolysin and neprilysin, and the metzincins MMPs and ADAMs (*46*, *47*), and C2H2 zinc fingers, where both motifs coordinate the ion with two histidines one helical turn apart (*48*).

**Figure 1:**
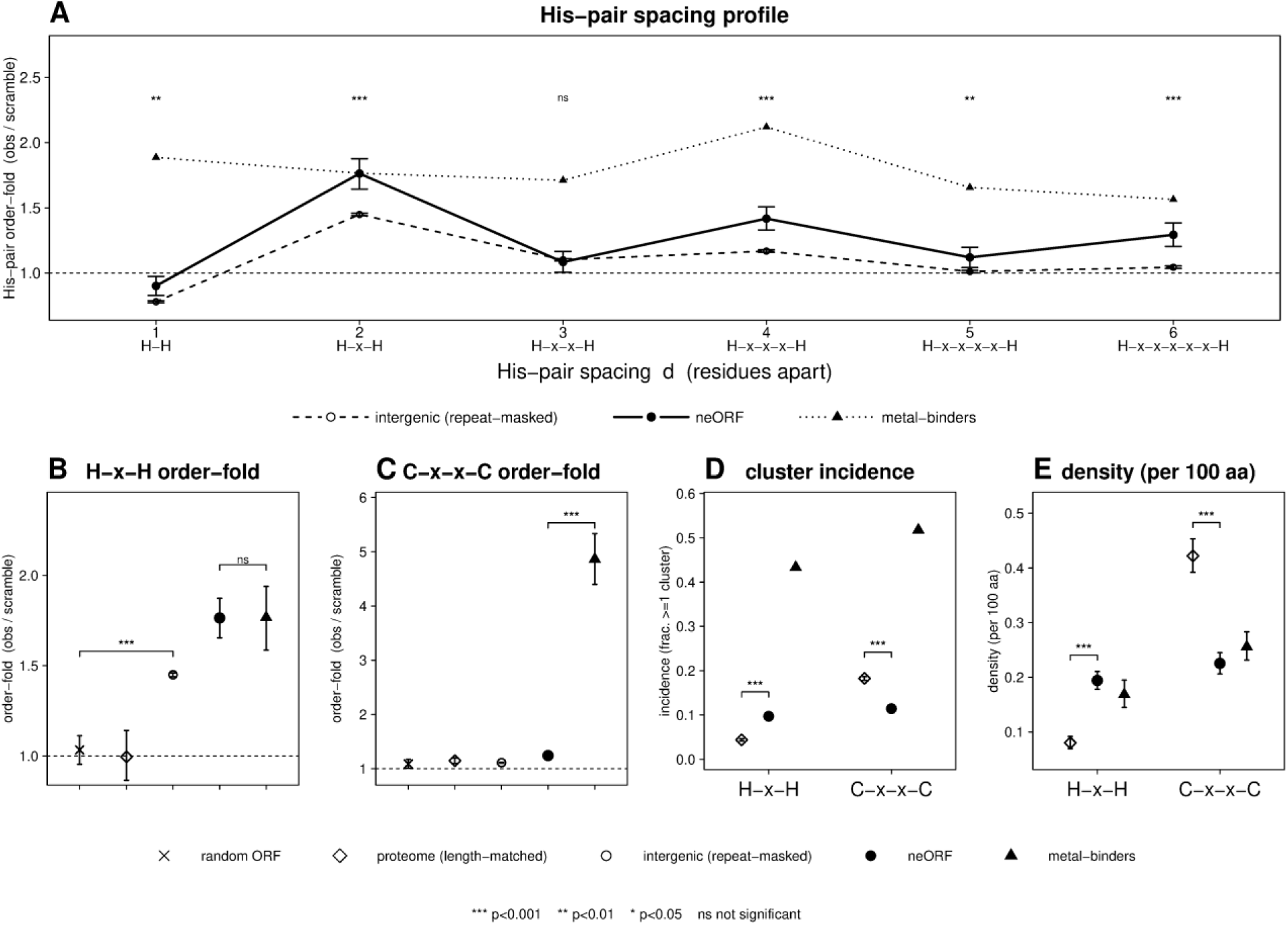
neORFs are enriched for H-x-H configurations but depleted of C-x-x-C configurations. (A) Relative number of H-(x)n-H configurations by gene class: neORFs (N=7,849), randomly selected intergenic, repeat-masked ORFs (N=1,472,446), and known metal-binding proteins (N=856). Raw numbers of configuration counts were divided by a bootstrapped distribution (k=3000) for each gene class. P-values were calculated between neORFs and a composition-matched null using Fisher’s Exact. Relative number of (B) H-x-H and (C) C-x-x-C configurations, and (D) cluster incidence and (E) density for H-x-H and C-x-x-C motifs in GC- and length-matched random ORFs, the length-matched *Drosophila* proteome, intergenic repeat-masked ORFs, neORFs, and known metal-binding proteins is shown.

Interestingly, repeat-masked intergenic sequences showed elevated levels of H-x-H sequences in comparison to GC- and length-controlled random sequences, suggesting that intergenic sequences may be poised for generating H-x-H sequences when compared to a permutation-based null that controls for composition (Figure 1B). One possibility is that such sequences arise from (CA)n dinucleotide repeats (see below). Additionally, another potential driving force may be recent pseudogenization of previously existing neORFs. The possibility of ancestral or residual protein-coding capacity is partially supported by the observation that the median length (*ℓ*_0.5_) of intergenic ORFs increases from 22 aa to 40 aa if the intergenic ORF has ≥ 2 histidine units. Similarly, the prevalence of at least 1 H-x-H motif increases in intergenic and neORFs as length increases. While some degree of increase is expected by chance alone, it is insufficient for explaining the progressive increase of H-x-H motif enrichment both in intergenic and neORFs (Table 1).

**Table 1:** H-x-H motif appearance in intergenic and neORFs by length. Observed H-x-H motif prevalence by length quartile in intergenic and neORFs progressively increases above an amino acid composition-matched shuffle performed within each quartile. P-values calculated using Fisher’s Exact. Uniformly low p-values for intergenic ORFs results from high N.

| intergenic ORFs |  |  |  |  |  |  |  |
| --- | --- | --- | --- | --- | --- | --- | --- |
| quartile | length<br>(aa) | median | ≥1 H-x-H | shuffle | obs/exp | p-value | His % |
| Q1 | 10–15 | 12 | 1.29% | 1.01% | 1.27 | <2x10<sup>-16</sup> | 3.10% |
| Q2 | 15–22 | 18 | 2.16% | 1.68% | 1.28 | <2x10<sup>-16</sup> | 3.15% |
| Q3 | 22–36 | 28 | 3.71% | 2.81% | 1.32 | <2x10<sup>-16</sup> | 3.23% |
| Q4 | 36–510 | 50 | 8.57% | 6.12% | 1.40 | <2x10<sup>-16</sup> | 3.37% |
| neORFs |  |  |  |  |  |  |  |
| quartile | length<br>(aa) | median | ≥1 H-x-H | shuffle | obs/exp | p-value | His % |
| Q1 | 30-45 | 37 | 3.66% | 2.99% | 1.23 | 0.29 | 2.81% |
| Q2 | 45-64 | 54 | 7.27% | 5.42% | 1.34 | 0.022 | 3.04% |
| Q3 | 64-92 | 75 | 10.63% | 7.29% | 1.46 | 2.7x10<sup>-4</sup> | 3.02% |
| Q4 | 92-619 | 117 | 17.34% | 12.01% | 1.44 | 2.6x10<sup>-6</sup> | 3.07% |

This latent potential for H-x-H motif appearance is also elevated when compared to a length-matched proteome sample (N=7,849, *ℓ*_0.5_=64 aa). The excess H-x-H found in both neORFs and intergenic ORFs in comparison to the length-matched proteome most likely reflects dilution/differences in amino-acid composition (Figure S1), though negative selection cannot be excluded. Interestingly, there is also no significant difference between neORFs and metal-binders specifically within H-x-H motifs (Figure 1B).

The preference for neORFs to show H-x-H motifs is striking in comparison to C-x-x-C chelation motifs. Here, random sequence, the length-matched proteome sample, intergenic sequence, and *de novo* neORFs appear to have no notable preference for C-x-x-C motifs. Alternatively, the C-x-x-C motif is a strong feature of known metal-binding proteins, showing a nearly 5x elevation of motif prevalence in comparison to all other categories (Figure 1C). This result suggests that the known class of metal-binding proteins have at least two distinct mechanisms by which they chelate metals, one that is shared with neORFs, and one exclusive to old, conserved metal-binding proteins (see “De novo genes as a potential zinc regulator”). An examination of both the raw incidence rate and the density of H-x-H vs C-x-x-C motifs further supports the conclusion that neORFs are preferentially enriched for bis-histidine coordinate chemistry, rather than C-x-x-C chelation. Specifically, while the incidence and density of these motifs in known metal-binders remains high, the length-matched proteome sample is depleted for H-x-H motifs in comparison to neORFs (incidence: neORF 9.7%, proteome 4.4%; Fisher’s Exact, p<2×10^-16^; density: neORF 0.194 motifs/100aa, proteome 0.080 motifs/100aa; Mann-Whitney, p<2×10^-16^), this pattern is reversed for C-x-x-C motifs (incidence: neORF 11.4%, proteome 18.3%; Fisher’s Exact, p<2×10^-16^; density: neORF 0.225 motifs/100aa, proteome 0.422 motifs/100aa; Mann-Whitney, p<2×10^-16^; Figure 1D-E). Interestingly, while the cluster incidence of H-x-H and C-x-x-C motifs is elevated for known metal-binding proteins in comparison to other gene classes, the density of these motifs is relatively low in metal-binding protein resulting from generally longer protein length (Figure 1E).

### (CA)n dinucleotide repeats drive potential neORF bis-histidine coordinate chemistry

While the consequences of H-x-H appearance are consistent with a selectable function via chelation (see *Discussion*), the H-x-H motif is not among the top 15 differentially prevalent SLiM present in the Eukaryotic Linear Motif database (*37*)(Figure S2). Additionally, while neORFs are enriched for histidine residues when compared to the length-matched proteome sample, neORFs are even more significantly enriched for cysteine than for histidine, thus providing an inadequate explanation for the simultaneous prevalence of the H-x-H but not the C-x-x-C motif (Figure S1). One possibility is that H-x-H motifs arise via the activation and lengthening of ORFs containing (CA)n repeats, the most highly abundant dinucleotide repeat found in the *D. melanogaster* genome (Table 2). Specifically, the sequence (CA)5 = CAC ACA CAC ANN, translates to His-Thr-His-(…). To test this hypothesis, I compared codon usage in the 2,036 histidines contained in H-x-H motifs to the 17,546 total histidines contained in neORFs, as histidine can be produced using either the CAC or CAT codon. This analysis revealed that histidines within H-x-H motifs were strongly biased towards utilizing the CAC codon over the CAT codon (56.2% vs 43.8%), consistent with (CA)n origination. This stands in contrast to codon usage across non-H-x-H neORF histidines, where codon usage was reversed (47.2% vs 52.8%)(Fisher’s Exact, p=3.0×10^-14^; Figure 2A). To further understand this phenomenon, I examined the relative frequency of amino acid usage for the middle position of the H-x-H motif. Consistent with the (CA)n hypothesis, I found that threonine was the most prevalent amino acid utilized in the middle position. When compared to background, threonine produced a 4.2x elevation over expectation (binomial test; p<2×10^-16^), suggesting that (CA)n dinucleotide repeats were the likeliest source of H-x-H motifs in neORFs. Note that while RepeatMasker was applied prior to later analyses, most (CA)n repeats from n=7 to n=9 typically are not masked due to their short length.

**Figure 2:**
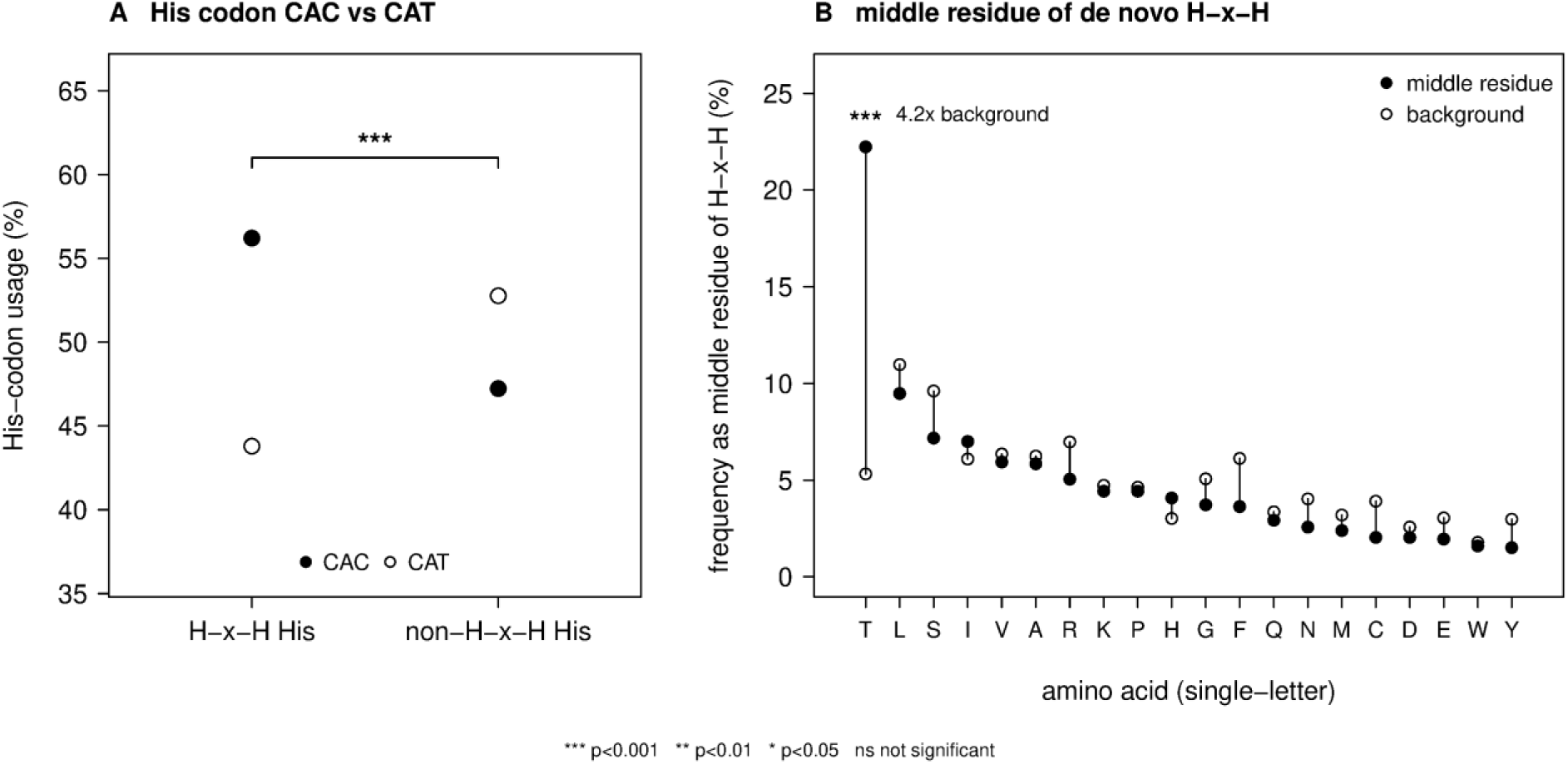
(CA)n dinucleotide repeats alter codon usage and amino acid positioning. (A) Histidine codon usage in both H-x-H motifs and non-H-x-H histidine residues found in *Drosophila* neORFs. Notably, histidines in H-x-H configurations are highly biased towards CAC usage. This trend is reversed when considering the set of non-H-x-H histidines found in neORFs. (B) Amino acid usage for middle residues (x) in bis-histidine motifs (H-x-H). The 4.2x elevation of Thr usage in the middle position is highly suggestive of (CA)n origins.

**Table 2:** Abundance of perfect dinucleotide microsatellite tracts by class in the *D. melanogaster* euchromatic genome. Perfect dinucleotide tracts (≥5 repeat units, ≥10 bp) were enumerated on chromosome arms 2L, 2R, 3L, and 3R of the UCSC dm6 assembly by our own reproducible census (see Methods; dinuc_census). Abundance is reported both by tract number and by total repeat length (bp). AC/(CA)n is the most abundant class by both metrics (∼1.6× the next class, AT), consistent with the AT-rich composition of the genome. Literature surveys of *Drosophila* microsatellites report the same qualitative dominance of AC/(CA)n.

| Dinucleotide class | Tracts (n) | % of tracts | Repeat length<br>(bp) | % by repeat bp |
| --- | --- | --- | --- | --- |
| AC / (CA)<sub>n</sub> | 11,323 | 52.9% | 159,936 | 54.0% |
| AG | 2,726 | 12.7% | 38,610 | 13.0% |
| AT | 7,271 | 34.0% | 96,830 | 32.7% |
| CG | 70 | 0.3% | 728 | 0.2% |
| *All classes* | *21,390* | *100%* | *296,104* | *100%* |

### A (CA)n-to-His-x-His spine converts microsatellite expansion into recurrent bis-histidine chemistry

The codon-usage and central-residue data point to a single, mechanistically explicit route from DNA to bis-histidine chemistry (Figure 3A). Read on the sense strand, (CA)n microsatellites present the repeating codons CAC·ACA·CAC·ACA…, translating as His-Thr-His-Thr…, where histidine residues fall at every second residue, with the intervening position fixed as threonine. Each additional CA unit that remains in frame thus appends one His-Thr pair and lengthens the run, whereas an out-of-frame unit shifts the reading frame and abolishes the motif, suggesting a single mechanism underlies the detected CAC-codon bias, the fourfold threonine enrichment at the central position, and the odd n-spacing (H-x-H, H-x-x-x-H) periodicity (Figures 1 and 2), without invoking selection at any individual locus.

**Figure 3:**
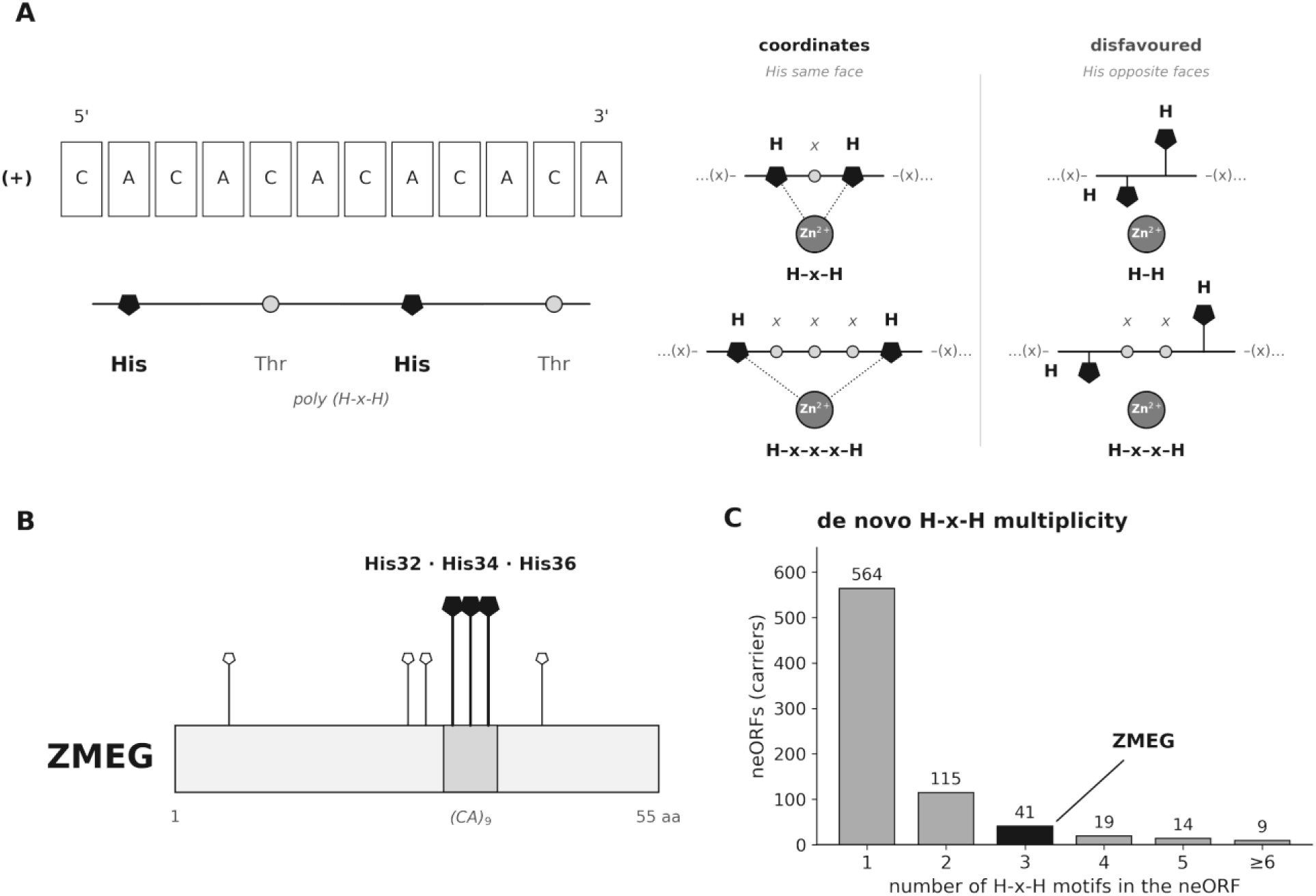
(CA)n microsatellite origins for ZMEG. (A) Illustration of (CA)n translation and cartoon of bis-histidine coordinate chemistry. (B) A genomic (CA)9 tract translates in frame to the His32/His34/His36 triple within a five-histidine run (His27–His36) contained in ZMEG. (C) H-x-H motif multiplicity across the uniquely translated neORF set (n = 7,849), with ZMEG indicated, demonstrates *de novo* proteins’ latent capacity for bis-histidine coordinate chemistry.

This pattern is readily apparent in the candidate chelation neORF *zmeygorynych* (ZMEG, Grandchamp designation UA_STRG.15519.1_1_193_357) (Figure 3B), which was identified as the most highly expressing poly-(H-x-H) neORF among the uniquely mappable, non-overlapping loci in a reanalysis of S2R+ expression data (see “Segregating neORFs are expressed in S2R+ cells”). Additionally, an analysis of ZMEG sequence shows that it originated by utilizing a His-Thr-His-Thr-His triple at positions His32/His34/His36 embedded within a five-histidine run (His27, His29, His32, His34, His36). Across the uniquely translated neORF set, 762 (9.7%) carry at least one H-x-H motif and 130 (1.7%) contain a poly-(H-x-H) His run of the kind produced by an in-frame (CA)n tract (Figure 3C). ZMEG lies in this poly-(H-x-H) tail. As a result, *zmeygorynych,* a 55-codon open reading frame on chromosome arm 3R, was named after the mythological Eastern Slavic dragon Zmey Gorynych and chosen to reflect a number of traits unique to ZMEG: 1) ZMEG was originally identified in an iso-female line of Ukrainian origin (*13*), 2) dragons are known for their propensity to hoard metals, 3) like ZMEG’s 3x H-x-H motif occurrences, Zmey Gorynych is typically described as possessing not one, but three heads, and 4) Zmey Gorynych’s autochthonous origins is reminiscent of ZMEG’s origination from ancestrally non-coding DNA.

### Translation of the complement of (CA)n

Translation of the complementary strand of (CA)n, (TG)n, appears as TGT·GTG·TGT… = Cys-Val-Cys, a C-x-C register. Unlike the even C-x-x-C motif, which may produce chelation activity on its own, the C-x-C motif cannot produce chelation by itself.

Because the neORF data set was derived from stranded transcriptome data (*13*), the strand from which each neORF is translated is known. It is therefore notable that (TG)n is present at near parity with (CA)n in both intergenic ORFs (CA:TG = 1.05, 51.1% CA; binomial test, p=1.9×10^-4^) and neORFs (CA:TG = 0.93; p=0.39), whereas (CA)n is favored in the whole proteome (CA:TG = 1.34; p=8.3×10⁻⁸). Accordingly, H-x-H and C-x-C enrichment each track their respective dinucleotide repeats. The two motifs are enriched to a similar degree over a permuted null in intergenic ORFs (1.45× and 1.39×; Fisher’s Exact, p<2×10^-16^ for both) and in neORFs (1.77× and 1.79×; p<2×10^-16^, p<2×10^-16^ respectively), whereas older genes are enriched 1.98-fold for H-x-H but only 1.13-fold for C-x-C (p<2×10^-16^, p=2.1×10^-6^ respectively; Figure S3).

Cysteine prevalence follows the same pattern, at 3.83% of residues in intergenic ORFs and 3.91% in neORFs (binomial test, p=7.3×10^-4^ relative to intergenic), but 2.25% in young genes (p<2×10^-16^) and 1.84% in older genes (p<2×10^-16^; Figure S3D). Histidine is an exception. Despite the increasing enrichment for H-x-H, histidine prevalence falls from 3.27% in intergenic ORFs to 3.02% in neORFs (p<2×10^-16^) and 2.42% in young genes (p<2×10^-16^) then finally 2.65% in older genes (p<2×10^-16^; Figure S3E), indicating that H-x-H enrichment reflects the spacing of histidine residues rather than their abundance.

The distinction between the odd C-x-C and even C-x-x-C spacings is reflected in known metal-binding proteins, which show no C-x-C enrichment (0.93×; Fisher’s exact test, p=0.38) but strong C-x-x-C enrichment (4.86×, p<2×10^-16^; Figure S3B, C), consistent with the C-x-x-C motifs found in zinc-binding folds (*49*). C-x-x-C enrichment is weak in intergenic ORFs (1.11×, p<2×10^-16^) and neORFs (1.24×, p=1.8×10^-7^), but rises to 1.78× in young genes and 1.93× in older genes (1.7×10^-34^, 3.5×10^-199^ respectively; Figure S3C).

Young genes provide an intermediate point between neORFs and established genes. Among Drosophila-restricted genes (including duplicate, *de novo*, and orphan genes)(*50*), C-x-C is not enriched above the expected baseline (1.01×; p=0.89), as is largely the case for older genes, whereas C-x-x-C is enriched 1.78× and H-x-H 1.37× (p<2×10^-16^, p=2.1×10^-9^ respectively; Figure S3A–C).

Together, these results suggest that the two chelation-competent registers, H-x-H and C-x-x-C, arise via different mutational routes. The H-x-H register is produced immediately and at high levels in the neORF pool via (CA)n dinucleotide repeats, while among established genes H-x-H spacing enrichment increases with gene age, from 1.37× in young genes to 1.98× in older genes (p=7.4×10^-11^). By contrast, (TG)n repeats yield only the non-competent C-x-C register, and C-x-x-C is weak in neORFs. Yet C-x-x-C is already present in young genes at near-ancestral levels (1.78× vs. 1.93×, p=0.13), so cysteine-based intra-motif chelation likely arises by routes other than dinucleotide repeats, whether through rapid acquisition after emergence or through gene duplication(*51*).

Despite the gradual transition from the C-x-C register to the C-x-x-C register in older genes, the significant enrichment of the C-x-C register over intergenic ORFs remains to be explained. Naively, the C-x-C register’s inability to chelate Zn^2+^ on its own, combined with a similar prevalence as the independently chelation-competent H-x-H register, could suggest that the preferential retention of both (CA)n and (TG)n dinucleotide repeats is a by-product of neutral processes, rather than selection for chelation in neORFs. However, it is possible that the observed enrichment for C-x-C motifs may not be directly related to the motif’s independent ability to chelate Zn^2+^ on its own, but in coordination with histidine or cysteine residues present elsewhere on the same protein unit or through multimeric interactions with other protein units (*Potential multimeric Zn^2+^ ion sequestration by de novo ORFs)*.

### Segregating neORFs are expressed in S2R+ cells

One necessary (but insufficient) prerequisite for presenting (CA)n microsatellites’ latent bis-histidine chemistry to selection is the transcription of these open reading frames in relevant tissue types. Given the association between innate immune response and Zn/Mn chemistry (see “De novo genes as a potential zinc regulator”), I therefore quantified neORF expression in *Drosophila* S2R+ cells using a publicly available RNA-seq series in which cells were exposed to control, zinc, and manganese conditions (GSE99332; *52*). Two quality-control steps proved essential. First, a subset of neORFs annotated as intergenic and non-overlapping in fact overlapped annotated features or repeats, or lay in low-mappability regions where multi-mapping reads inflate apparent expression. Second, restriction to uniquely mapping reads and to loci passing a per-base mappability filter yielded 472 high-confidence, uniquely mappable, non-overlapping *de novo* ORFs (Materials and Methods)(Figure 4A, B).

**Figure 4:**
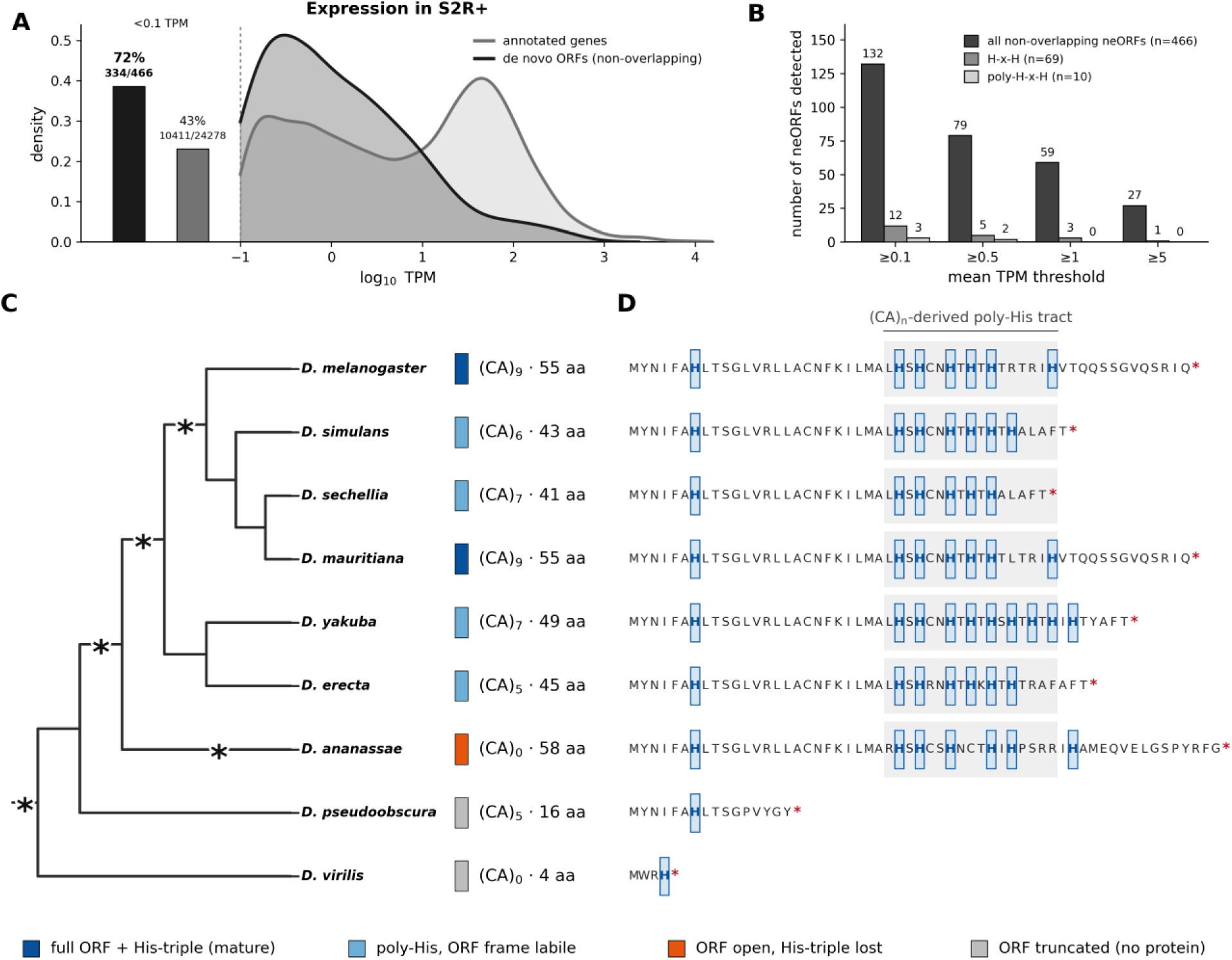
*De novo* neORF transcription and the origins of ZMEG. (A) Expression (mean TPM across all conditions) of uniquely mappable, non-overlapping *de novo* ORFs (n = 466) versus annotated genes in S2R+ cells. Left bars provide the fraction of undetected transcripts (<0.1 TPM). (B) Number of *de novo* ORFs transcribed at increasing mean-TPM thresholds for all neORFs and for the H-x-H (n = 69) and poly-(H-x-H) (n = 10) subsets. (C) *De novo* origin and activation trajectory of ZMEG across nine *Drosophila* species (branch lengths not to scale), including (CA)n tract length, ORF length to the first stop, and His-motif status mark. Asterisks indicate polarized evolutionary events described in the main text. (D) ORF translations (ATG to first stop) aligned across all nine species: histidines are boxed and (CA)n-derived poly-histidine tracts are shaded.

This analysis revealed that *de novo* ORFs are transcribed, but at characteristically low levels (*13*). 28% (132/466) of these neORFs reached a mean expression of ≥0.1 TPM (transcripts per million), compared with 57% of annotated genes, while 72% of neORFs were effectively undetected (<0.1 TPM) (Figure 4A). Only a small fraction of neORFs exceeded 1 TPM (Figure 4B). Expression was nonetheless constitutive rather than sporadic, as detected neORFs were expressed across replicates and conditions. Thus, the low absolute levels reflect weak, pervasive transcription rather than tight, condition-specific regulation. Notably, the most highly expressed neORFs were not H-x-H carriers (Dataset S1).

Interestingly, ZMEG is expressed at low levels (mean ≈0.9 TPM across conditions, ≈0.5 TPM in controls) and rises under manganese (≈1.5 TPM), an apparent ∼2.8-fold increase (DESeq2 maximum-likelihood log_2_ fold-change ≈ +1.5). Intriguingly, a second *de novo* neORF bearing an H-x-H motif was expressed at appreciably higher levels than ZMEG (≈4.8 TPM; Dataset S1), but lacked the signature H-T-H motif. Regardless, this rise is not statistically significant. In the manganese-versus-control comparison, ZMEG has an adjusted p=0.24 (the same comparison underlying the +1.5 log2 fold-change above), and it is likewise non-significant versus zinc (p=0.79; Materials and Methods). More importantly, manganese is a broad, strong stressor in these data: 3,900 genes are differentially expressed under manganese versus 859 under zinc (roughly 4.5-fold more), with classic proteotoxic and metal-detoxification programs (Hsp70, metallothioneins) strongly induced. ZMEG’s modest rise is therefore best read as part of a general stress-mobilization wave that broadens the pool of transcribed cryptic and *de novo* loci, not as a dedicated manganese response (see Discussion).

Consistent with cryptic rather than dedicated transcription, ZMEG’s promoter-proximal region contains no canonical core promoter (no TATA box) and shows no enrichment of MTF-1 metal-response elements (MREs) above intergenic background, in contrast to the bona fide MTF-1 targets MtnA–E, whose promoters are strongly MRE-enriched (Materials and Methods). ZMEG is thus not an MTF-1/MRE-regulated locus, reinforcing the interpretation that its expression, along with its manganese-associated increase, reflects pervasive, stress-amplified transcription rather than a locus-specific metal-response circuit.

In order to confirm that ZMEG’s expressed sequence is genuine and matches its assembly, I reconstructed the ORF directly from the S2R+ reads, rather than from the reference alignment. This analysis recovered a sequence identical to the Grandchamp et al. (*13*) assembly, including the in-frame (CA)9 tract and the His32/34/36 triple. Additionally, because dinucleotide tracts are prone to alignment artifacts, tract length and reading frame were read by eye from the raw reads for ZMEG and its comparators throughout (Materials and Methods).

To determine whether ZMEG is representative or exceptional, I compared the neORF’s two key properties ((CA)n-derived poly-(H-x-H) signature and stress-associated expression) across the *de novo* set, revealing a decoupling of these two properties. Indeed, the latent (CA)n-to-H-x-H reservoir is genomically pervasive. In the full uniquely translated neORF set (n = 7,849), 762 loci (9.7%) carry at least one H-x-H motif and 130 (1.7%) poly-(H-x-H) runs (Figure 3C). Expression, however, remains limiting, as further analysis of the 466 uniquely mappable, non-overlapping neORFs quantified in S2R+ cells shows that 69 carry an H-x-H motif and 10 poly-(H-x-H) runs (Figure 4B). Of these ten poly-(H-x-H) loci, three are detectably transcribed, ZMEG, UA_STRG.5397.1, and UA_STRG.826.1, with ZMEG the most highly expressed and the only one showing metal-induced upregulation (Dataset S1), suggesting that transcription serves as a significant barrier for the activation of individual neORFs.

### ZMEG arose de novo in the melanogaster lineage from conserved ancestral sequence

To validate ZMEG’s *de novo* origins and to reconstruct how its metal-relevant chemistry arose, the locus was traced across nine *Drosophila* species spanning roughly 50 My of divergence (Materials and Methods)(Figure 4C,D, Table S1). Critically, the ZMEG locus is syntenic in all nine genomes, where the region lies in the same intergenic interval flanked by the conserved single-copy anchors Gr93 and glec, and the upstream sequence is largely shared across lineages. This observation confirms ZMEG’s in-situ origination, positively excluding the possibility that ZMEG is an orphan gene.

This analysis also revealed that the ATG and the immediately following N-terminal codons are conserved across the *melanogaster* group and into *D. pseudoobscura* (∼25 My), which shares ZMEG’s N-terminal peptide (MYNIFAHLTSG…). However, this ORF terminates after 16 codons upstream of the (CA)n tract. Importantly, the syntenic locus itself is recoverable in the most distant outgroup, *D. virilis* (∼40–50 My), establishing a deep nucleotide-level anchor. However, *D. virilis* translates only a short, divergent N-terminal peptide and encodes no ancestral ZMEG protein. These results confirm that the full-length ORF and its bis-histidine chemistry are derived solely within the *melanogaster* lineage in concordance with its identification as a segregating neORF (*13*), and was not inherited from an ancient protein.

Critically, polarizing character states on the tree resolves a stepwise, microsatellite-driven birth mechanism for ZMEG (Figure 4C). In the two most distant outgroups the ancestral ZMEG reading frame terminates before the (CA)n tract. On the *melanogaster*-group stem, the ancestral ZMEG reading frame extends past this ancestral early stop codon, opening and elongating the nascent ORF. Interestingly, an in-frame (CA)n tract seeds a poly-histidine motif within the *melanogaster* sub-group. Finally, the tract reaches (CA)9 in *D. melanogaster* and *D. mauritiana*, translating to the full poly-(H-x-H) run with the His32/34/36 triple in a mature 55-codon ORF. Crucially, this mature state is tract-length-dependent and hypervariable in *D. simulans* ((CA)6, 43 codons) and *D. sechellia* ((CA)7, 41 codons), as the same complex carries poly-histidine as a shorter, frame-labile ORF. Alternatively, *D. yakuba* and *D. erecta* carry poly-histidine at intermediate tract lengths.

The same lability is evident within *D. melanogaster*, where the tract segregates between 6 and 10 units across iso-female strains studied in (*13*), with in-frame deletions and frameshifting indels that lengthen, shorten, or disrupt the His run (Table S1). The (CA)n tract is thus a reservoir in flux rather than a fixed feature. Notably, ZMEG arises directly in-frame, with no clear evidence of an intervening functional-RNA stage, suggesting that its origin did not proceed through a cultivated ancestral noncoding-RNA intermediate (*19*).

### ZMEG’s His32–His36 pair is a candidate bis-histidine metal site

Three independent lines of evidence converge on a single candidate site formed by His32 and His36 (Figure 5). First, the above joint evolutionary-mechanistic analysis indicated that the His32/34/36 triple is the direct in-frame translation product of the (CA)9 tract, so the site sits exactly where the (CA)n spine predicts bis-histidine chemistry should appear. Second, geometry: an analysis of ZMEG’s ESMFold-predicted structure (Materials and Methods)(*53*) models the His32 and His36 side chains residing 4.63 Å apart (imidazole-nitrogen separation), within the range spanned by two-histidine metal sites, and a modeled Zn^2+^ sits exactly at 2.31 Å from each coordinating nitrogen (Figure 5A). Third, a learned zinc-site predictor, Metal3D (Materials and Methods)(*54*), places its highest-probability voxel for ZMEG just 0.8 Å from the midpoint of the His32–His36 pair (Figure 5B). While such structure predictions do not fully establish the existence of metal bonding, the convergence of two structural predictors operating on the predicted fold, together with the independent evolutionary argument, yields a specific and falsifiable prediction: that ZMEG’s putative chelating function lies in the His32–His36 pair’s capacity for bis-histidine coordinate chemistry. Notably, unlike the idealized β-strand register of the (CA)n schematic (Figure 3A), ZMEG’s predicted fold is helical, and the coordinating histidines are related by an i,i+4 spacing (His32→His36) consistent with same-face presentation on a helix (Figure 5A,C). The intervening residue at ZMEG’s His-x-His positions is threonine, within the substrate tolerance of the histidine methyltransferase METTL9. This raises the possibility of a post-translational “off” switch (see *De novo genes as a potential zinc regulator*).

**Figure 5:**
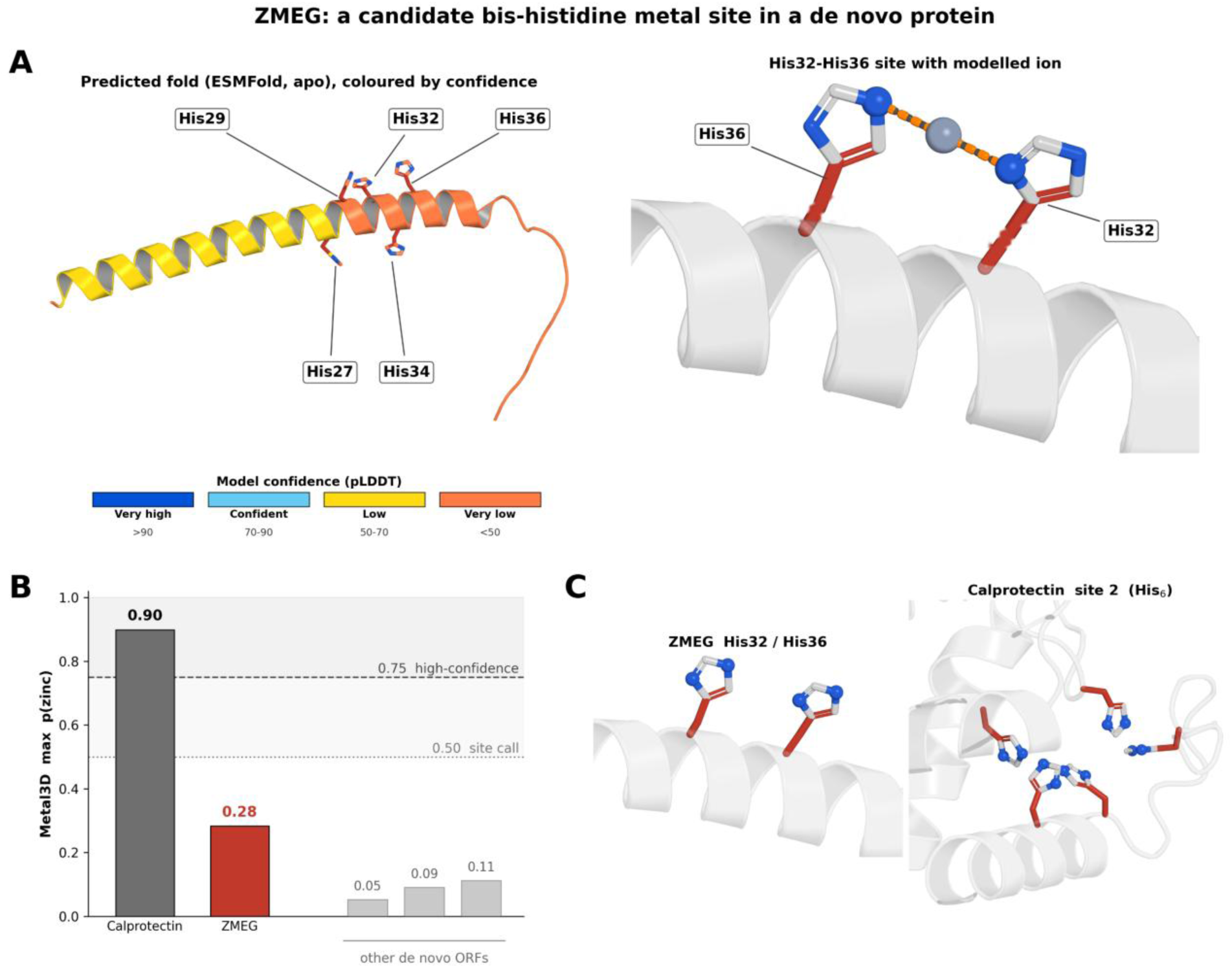
A candidate bis-histidine metal site in ZMEG. (A) ZMEG predicted fold (ESMFold, apo; colored by pLDDT model confidence) with the His27/29/32/34/36 run indicated, and a close-up of the His32–His36 site with an illustrative modeled Zn^2+^ (apo prediction; imidazole-nitrogen separation 4.63 Å, metal–nitrogen 2.31 Å). (B) Maximum predicted zinc-site probability (Metal3D): ZMEG (0.28) versus a calprotectin S100A8/S100A9 positive control (0.90) and other *de novo* ORFs (0.05–0.11); dashed lines mark the tool’s 0.5 and 0.75 confidence thresholds. For ZMEG the highest-probability voxel lies 0.8 Å from the His32– His36 midpoint. (C) Structural comparison of ZMEG’s loose two-histidine pair with calprotectin’s six-histidine metal cage.

### Potential multimeric Zn^2+^ ion sequestration by de novo ORFs

Given the chelating capacity of both cysteine (via its thiol group) and histidine (via its imidazole group), one possibility remains that cysteines and histidines work together, either through intra- or inter-molecular interactions to form a single chelating unit. A notable exemplar of this possibility is the dedicated Zn^2+^ chelator calprotectin in mammals which functions as a heterodimer (Figure 5)(*55*). To test whether neORFs may be chelating free Zn^2+^ ions through a multimeric state, I arranged two copies of ZMEG into the homodimeric configuration that it would require to bind Zn^2+^ (Figure 6). This configuration produced idealized tetrameric binding with Zn^2+^, boosting the monomeric Metal3D score from 0.28 to the homodimeric score of 0.95, a value on par with calprotectin’s 0.90 Metal3D score. Notably, this configuration also avoids residue overlap in 3D space, which would disfavor such binding due to steric hindrance or entropic effects. Critically, such a dimeric configuration also decreases the probability of random Zn^2+^ binding to free histidine or cysteine residues across the entire proteome, thus reducing the risk of Zn^2+^-bridged protein aggregation. This observation also raises the possibility that the entire class of chelation-competent neORFs may be acting in concert to sequester free Zn^2+^ ions.

**Figure 6:**
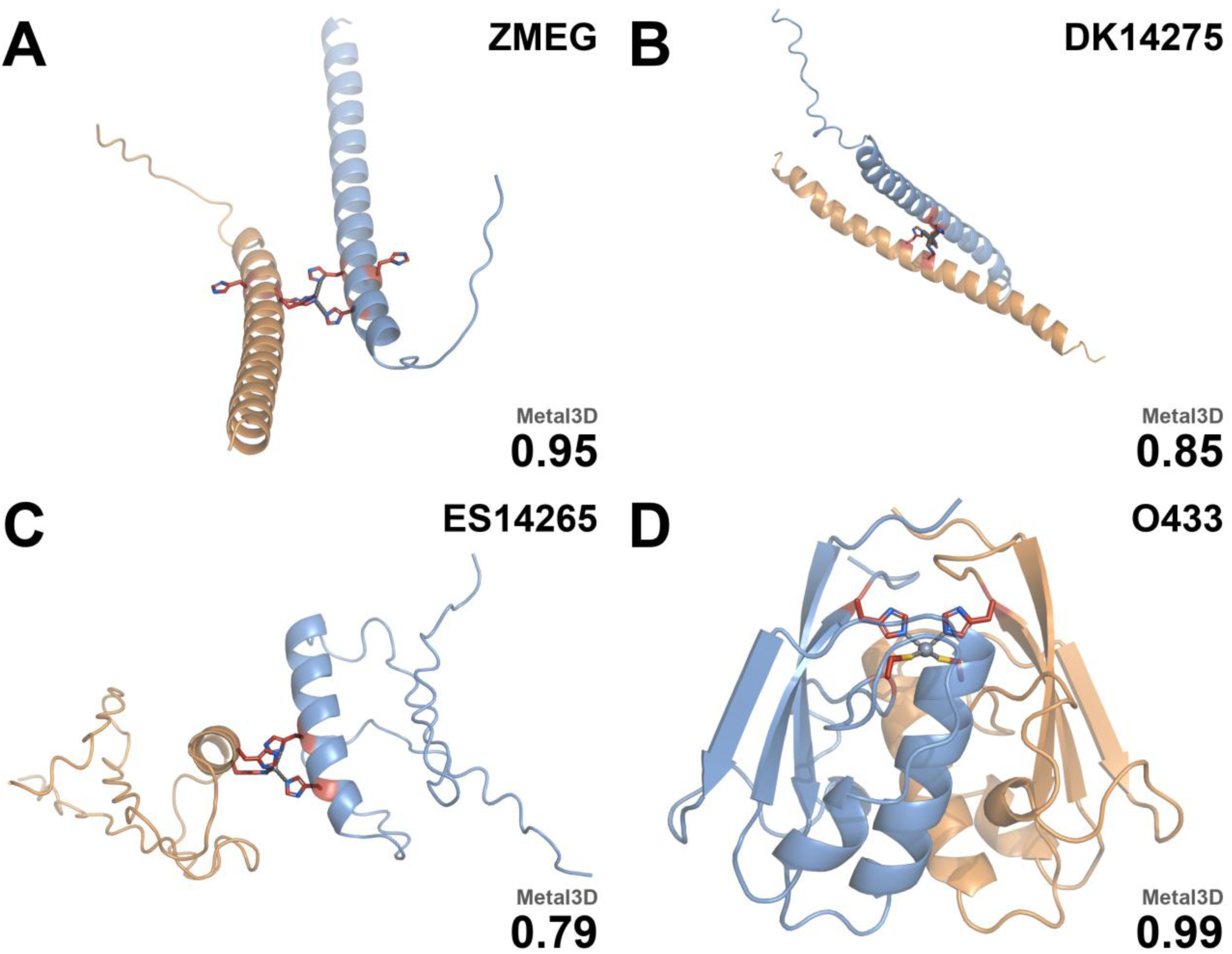
Dimeric configurations of neORFs. Predicted Zn^2+^-bound dimeric structures and associated Metal3D scores are shown for (A) ZMEG, (B) DK14275, (C) ES14265, and (D) O433.

In order to more systematically test for this possibility, I took a number of neORFs that produced relatively high Metal3D scores and predicted possible Zn^2+^-binding homodimeric configurations using Boltz-2, a machine learning model designed to predict small molecular-protein binding interactions (Materials and Methods)(*56*). The resulting configurations were then scored using Metal3D, demonstrating the latent capacity for neORFs to bind free Zn^2+^ in multimeric configurations (Figure 6). An analysis using a random sample of potential pair-wise interactions showed similar results.

One recurring and highly-evolved feature of dedicated Zn^2+^ protein interaction partners, such as zinc-finger nucleases or chelators like calprotectin, is the prevention of Zn^2+^-bridged protein aggregation via ionic sequestration within cage-like structures. My tests of homodimeric Zn^2+^ binding revealed that neORFs also possess a latent capability for producing similar cages. In particular, one still-segregating neORF, which I refer to as ORF433 (O433, Grandchamp designation FI_STRG.14643.1_8_2494_2775), forms a Cys2-His2 cage surrounding free Zn^2+^ ions with a Metal3D score of 0.999. This open reading frame is named after the famous work by John Cage, 4’33”, which challenged contemporaneous paradigms of music by transforming ambient noise into the performance itself.

Interestingly, an analysis of O433’s origination reveals how the (TG)n repeats, not (CA)n, generates O433’s putative Zn^2+^ binding function between Cys62 and His89. Cys62 is produced as a direct result of an existing (TG)n tract. This tract exists with n=9 in the dm6 allele, while O433 contains n=10. This addition of an extra TG dinucleotide induced a frameshift downstream of Cys62 which thus generated His89. Notably, both O433 and the dm6 allele contain a (CA)n tract that forms a H-x-H-x-H motif on histidines His5, His7, and His9 which generate a Metal3D score of 0.48 alone. It is unlikely that the homodimeric dm6 allele can sequester Zn^2+^ alone, as 1) Boltz-2 does not predict such a configuration and 2) an idealized Zn^2+^ binding configuration is not possible due to steric hindrance, thus likely requiring an interaction partner that could potentially take the form of a different ORF such as ZMEG. This observation provides a potential solution to Zn^2+^-bridged aggregate formation in a two-fold manner, both through direct sequestration of Zn^2+^ within a cage-like structure while also allowing for the termination of putative outward-facing Zn^2+^-bridged binding via an ORF like ZMEG. Such improvement over the functioning of the dm6 allele could potentially provide a selective effect that is large enough to overcomes the drift barrier resulting from the Dykhuizen-Hartl effect (*De novo genes as an evolvable plasticity mechanism,* Supplementary Text) (*57*, *58*).

## Discussion

This work connects, for the first time, a single *de novo* protein-coding gene with an explicitly connected evolutionary and molecular mechanism. By bridging *de novo* gene origination, zinc handling in *Drosophila*, and concrete biochemistry, this work introduces the possibility of broader generalization of ligand-binding activity as a driver of *de novo* protein origination. Importantly, the evolutionary (CA)n-to-H-x-H axis makes this connection tractable because the same microsatellite that can be polarized on a phylogeny also specifies, through the reading frame, the chemistry of the resulting peptide, so that evolutionary history and candidate function are encoded in a single feature.

### De novo genes as a potential zinc regulator

METTL9 is a methyltransferase that has been recently demonstrated to play a key part in post-translational regulation of a broad range of substrate proteins via pervasive 1-methylhistidine modification (*44*). METTL9 tolerates only a limited set of residues at the intervening (x) position, including threonine, which is the most common residue for (x) found in this analysis. As METTL9 does not specifically prefer threonine over other members of the A-N-G-S-T motif (*44*), the enrichment of threonine in particular reflects (CA)n origins rather than co-adaptation to METTL9. Notably, mammalian METTL9 methylates a histidine within the H-x-H motif to install Nπ 1-methylhistidine, which acts as a post-translational off-switch by decreasing the metal-binding capability of target proteins. This has been shown to be the mechanism for calprotectin, the mammalian anti-bacterial S100A8/S100A9 heterodimer, of which the short, disordered S100A9 (MRP14) subunit is the methylation target (Figure 5C)(*59*). Structurally, calprotectin’s manganese/zinc “site 2” is a hexahistidine cage assembled across both subunits (S100A8 His17/His27 and S100A9 His91/His95/His103/His105)(*55*), whose six histidine donors far exceed ZMEG’s single candidate two-histidine pair. While calprotectin is conserved between humans and mice (*60*), no S100/calprotectin orthologue is annotated in *Drosophila*. Nπ-methylation of S100A9 His107 weakens its zinc affinity roughly sixfold (the dissociation constant rises from ∼14.3 to ∼90.9 µM) (*45*). Notably, upon *Staphylococcus aureus* infection, the methylation of S100A9 His107 is reduced (*45*). This loss of methylation thus increases local sequestration of zinc, which starves *S. aureus* of metal ions and clears the infection. In line with this, METTL9-KO mice with constitutively unmethylated S100A9 were found to be more resistant to *S. aureus* infection, consistent with calprotectin’s zinc-sequestration function as the causal mechanism of resistance (*45*).

In *Drosophila*, the METTL9 ortholog (*Mettl9*/*CG5339*) is also an Nπ 1-methylhistidine His-methyltransferase, though this regiochemistry is inferred by orthology and has not been directly confirmed for the fly enzyme. Importantly, *D. melanogaster* has a known metal response pathway, where copper, cadmium, silver, and zinc ingestion upregulates the MTF-1 transcriptional pathway (*61*, *62*) that regulates known copper transporters, e.g. Ctr1B and ATP7 (*63*), and known metallothioneins, e.g. MtnA–E (*64*). However, these metallothionein proteins are cysteine-rich, not histidine-rich. Additionally, unlike copper, cadmium, and silver, zinc only weakly induces the metallothionein arm of the MTF-1 response, whereas the transporter arm, e.g. ZnT35C, is robustly zinc-induced and MTF-1-dependent (*65*).

On the other hand, zinc regulation in *D. melanogaster* remains relatively understudied (*65*), where “little is still known about local redistribution and availability of Zn within… the cell.” (*66*). What is currently known about Zn response in fruit flies is mostly limited to zinc transporters such as ZnT35C (*65*), not direct zinc chelators. Concurrently, while zinc sequestration is well-characterized in the previously reported mouse infection model, S100A9/calprotectin is a vertebrate-specific innovation as previously noted, with no identifiable homolog in *Drosophila*. However, previous work has noted that fruit flies nonetheless mount a Transferrin-1–mediated nutritional-immune response during infection (*67*). As of publication, there are currently no known dedicated histidine-rich zinc-sequestration or buffering proteins in *Drosophila*, suggesting that this key biological function may be served by the collective action of an entire family of mutually unrelated, strain-specific neORF zinc chelators, rather than a more traditionally ascribed duplication-based expansion of a known gene family.

Interestingly, recent work has demonstrated that *Drosophila Mettl9*/*CG5339* has altered binding affinities for known human METTL9 targets. Combined with S100A9/calprotectin’s known *zinc*-binding function in mammals, and METTL9’s known mechanism of H-x-H methylation, my results suggest that *D. melanogaster*’s zinc response may potentially be acting through a separate, yet undercharacterized pathway, via the entire *class* of *de novo* H-x-H proteins. Such interactions would be consistent with the documented *Drosophila*-specific evolution of *Mettl9* binding, where substitutions in flanking and -x-residues were found to be less tolerated than with human METTL9 (*68*). In that study, the authors demonstrated how *Drosophila Mettl9* had increased stringency in activity, preferring to bind A, G, S, C, or T residues, rather than N, Q, H, and P in the middle position. ZMEG provides a concrete candidate for this logic: its central (x) residue is threonine, which lies within the more stringent *D. melanogaster* METTL9 acceptor set (A/G/S/C/T; *68*), so Nπ-methylation of a histidine in its H-x-H motif would offer a plausible post-translational off-switch on a candidate *de novo* chelator, mirroring METTL9’s demonstrated control of calprotectin’s zinc affinity (*45*).

### Promiscuous expression presents de novo H-x-H motifs to selection

For the latent metal-coordinating chemistry of a *de novo* H-x-H peptide to be visible to natural selection, its open reading frame must be transcribed and translated. Notably, however, neither step requires locus-specific regulation. Pervasive, low-level transcription of intergenic sequence is a general feature of eukaryotic genomes (*69*) and is the very property by which the neORFs analyzed here were identified as transcribed loci (*13*). Noncanonical and short ORFs are similarly engaged by ribosomes at appreciable rates (*70*, *71*). This promiscuous transcription and translation are expected to intensify under physiological stress as transcriptional and translational specificity relaxes. For example, previous work has shown that translational fidelity and start-site selection loosen during metal stress (*72*). Under such conditions, the pool of expressed *de novo* ORFs transiently broadens, sampling from a large reservoir of (CA)n-derived H-x-H peptides without any single locus having evolved a dedicated promoter. Consistent with this, *de novo* genes tend to originate in ancestrally closed chromatin and acquire expression through newly originated regulatory sequences rather than by co-opting preexisting open chromatin (*73*); pervasive, stress-amplified transcription thus offers a route by which such initially unregulated loci, including those bearing (CA)n-derived H-x-H motifs, can be presented to selection before dedicated de-repression and/or heterochromatin escape evolves.

This stress response supplies the missing bridge between motif supply and selectable function. Specifically, (CA)n expansion can generate the latent capacity for H-x-H motif emergence in neORFs. Subsequently, promiscuous, stress-amplified expression then presents these nascent peptides and their bis-histidine coordinate chemistry to selection, which can retain and quantitatively tune those whose metal-coordinating activity proves beneficial. Critically, this route requires no sequence-specific regulatory innovation at any individual locus: a general, non-specific expression mechanism suffices to expose a compositional property (histidine spacing) to selection. This parallels the adaptive, evolutionarily tuned phase behavior of stress-responsive proteins, whose selectable function is a bulk sequence property realized without site-specific regulation (*74*, *75*), and the broader pre-adaptation view in which function can precede and drive the establishment of *de novo* genes (*14*).

Because it is a general property rather than a locus-specific adaptation, the transcriptional signature of this mechanism is expected to be diffuse and non-motif-specific. This is consistent with, though not resolved by, existing metal-challenge transcriptomic studies (see “Segregating neORFs are expressed in S2R+ cells”), and would be most directly testable by ribosome profiling under acute metal stress. However, it should be noted that pervasive, non-specific low levels of translation are abundant as well. As such, ribosome profiling still would not provide a “smoking gun” demonstration of the chelation hypothesis, as observed levels of non-specific translation would remain indistinguishable from noise. Consistent with this general, non-specific reading, the metal-challenge data analyzed here show a strongly asymmetric response, as manganese, a broad stressor, differentially regulates roughly 4.5-fold more genes than zinc (3,900 versus 859). Here, ZMEG’s own manganese increase is not individually significant (adjusted p=0.24), consistent with the diffuse, non-locus-specific signature this model predicts (Dataset S1).

### De novo genes as an evolvable plasticity mechanism

Emergence bias is a critical feature underlying the dynamics of genome evolution (*25*). Specifically, natural selection can only select for features that are presented to it *in the order of emergence*. That is to say, natural selection cannot select for variants that have not yet appeared in a population. In species with high effective population sizes such as *D. melanogaster*, this evolvability problem is amplified, as slightly deleterious variants are more effectively purged in such populations, preventing “tunneling” behavior between selectively beneficial states (*76*). This crucial detail is a defining aspect that separates the study of evolutionary genetics and the study of random peptides and/or directed evolution.

Notably, the same efficient selection has a complementary consequence for the types of functional but redundant reservoirs proposed in this work, an effect previously described as the Dykhuizen-Hartl Effect (*57*, *58*). As aggregate buffering capacity is supplied across many loci and saturates demand, the marginal selection coefficient at any single locus likely falls below the drift barrier (*77*) even at *D. melanogaster*’s effective size (*78*, *79*), so no per-locus selective signature is expected (Supplementary Text). This does not, however, exclude the possibility of large-effect loci from emerging.

The emergence of H-x-H motifs, potentially in connection with the prevalence of (CA)n dinucleotides in *Drosophila*, offers the tantalizing possibility that a subset of *de novo* proteins have independently but collectively evolved to act in a module-like manner. Such collective transcription and translation could allow for the potential quantitative tuning of zinc response via alteration of the mutational target size of plastic response to metal consumption or bacterial infection. Such mutational target size, and thus the relative mutation rate of genic vs epigenetic responses to environmental shifts, has been shown to play a role in determining the long-term fate of plasticity (*80*). Notably, AC/(CA)n is the most abundant class of dinucleotide microsatellite in *Drosophila melanogaster*, accounting for ∼53% of perfect dinucleotide microsatellite tracts (≥5 repeat units) on the euchromatic chromosome arms and ∼54% by repeat length (Table 2), despite the genome’s overall AT-richness (AT tracts form a substantial second class at ∼34%). Most (CA)n tracts are short (median 6 units), but a minority extend into a longer tail; tracts such as ZMEG’s (CA)9 sit at this chelation-competent extreme in *Drosophila* (*81–84*), yielding approximately 2-3 His-Thr repeats. This well-documented prevalence of (CA)n microsatellite repeats in *Drosophila* thus produces His-Thr-His upon translation of the (+) strand, and Cys-Val-Cys (but not the chemically active Cys-Cys or Cys-x-x-Cys motifs) on the (-) strand . While the evolutionary forces underlying the limited appearance of longer (CA)n tracts remain a mystery (*85*), one outcome of these repeats is the likely rapid appearance of a wide class of intrinsically disordered proteins with the ability to sequester metal, related not by direct descent via duplication, but the repeat, independent origination of hundreds or thousands of the same protein *subunit* – again, arising *de novo* and not through duplication.

The possibility of such independent origination highlights the key role of evolvability in natural selection – this a mechanism is plausible only when such events are heavily biased to emerge (*25*). Similar effects arising from functional repeat units accelerating the evolution of genic solutions to fluctuating environments have been previously observed in bacterial promoters (*86*). As a dinucleotide repeat, it is important to note that expansion of this highly labile repeat is also likely to produce frameshifts within open reading frames, causing the H-x-H motif to be both highly evolvable but similarly easy to lose. As the mutation rate of such dinucleotide repeats is much higher than more commonly studied single nucleotide substitutions (*87–89*), this may allow H-x-H motifs from (CA)n repeats to potentially serve a secondary function as a favored launchpad for generating further protein coding diversity.

### A latent reservoir and the limits of a genomic lens

Two important limitations must be acknowledged in framing this study. First, the metal-challenge series analyzed here may not reach the concentrations, duration, or cellular context of a physiologically relevant metal-stress episode, so the absence of a strong, ZMEG-specific transcriptional response in these data does not bound the reservoir’s potential. Instead, it may simply indicate that the relevant regime was never sampled. Second, ZMEG is expressed at levels that would be easily missed without targeted analysis. Consistent with this, a companion population-level survey of the same neORF catalog (*90*) detected roughly half of these neORFs as expressed across adult tissues, developmental stages, sexes, and African and European populations. As a segregating ORF, this includes low-level, broad expression of ZMEG (OG0004162) within specific lineages. Notwithstanding ZMEG’s original identification in (*13*), such expression became visible when expression was assayed in a defined cell population (S2R+ cells). The capacity for (CA)n-derived bis-histidine chemistry may sit in a latent, genomic reservoir (*91*), as demonstrated through a single ORF (ZMEG) that had previously gone unnoticed in published data sets. Here, I have shown that ZMEG’s conversion into a transcribed, candidate metal-binding peptide is not merely feasible but, given the genomic prevalence of the underlying microsatellite, plausibly recurrent.

This same genomic reservoir is, paradoxically, extraordinarily difficult to interrogate through a genomic lens. If the mechanism proposed herein is a general, stress-amplified broadening of expression rather than a locus-specific circuit, then its signature in transcriptomic and/or translatomic data is expected to be indistinguishable from noise. Accordingly, sequencing cannot by itself distinguish a functionally engaged *de novo* chelator from incidental pervasive expression/translation. The two questions that would settle function are not genomic at all: does ZMEG actually bind metal inside the cell? and does genetically deactivating the (CA)n reservoir measurably shift intracellular metal levels? Neither is answerable by sequencing alone, highlighting the need for further experimentation. Beyond just a mere caveat, this work highlights, rather than weakens, the value of this work: the entire evolutionary-to-molecular account assembled here, including origin, mechanism, and a testable candidate site, was recovered from a handful of publicly available datasets (a single focal species, eight outgroup genomes, and one previously published RNA-seq dataset), for a gene effectively invisible to standard genomic pipelines. Crucially, a motif-first search also would not have found ZMEG: one would have had to known the yet unannotated (ELM database)(Figure S2) H-x-H signature in advance, whereas here, the evolutionary analysis drove and surfaced the underlying chemistry. I highlight that this inversion is the broader thesis this work is meant to advance, demonstrating that comparative and population-level study of how sequences arise and change can serve not only as a post-hoc historical description of molecular history, but as a discovery tool for uncovering new biochemistry and biophysics.

## Materials and Methods

### New expressed open reading frame (neORF) dataset

7,894 neORFs from Grandchamp et al. (*13*) were retrieved from the associated data archive (Zenodo record 7322757). Because the published protein sequences appended stop-codon-derived nucleotides as C-terminal residues, neORFs were re-translated from their nucleotide entries in frame, terminating at the first in-frame stop. Re-translations were de-duplicated by amino-acid sequence, yielding 7,849 unique *D. melanogaster*-specific neORFs (median length ℓ₀.₅ = 64 aa). Following Grandchamp et al., neORFs are the *de novo* ORFs segregating in fewer than 7 of 7 independent iso-female lines, where neORFs are young, transcriptionally supported, population-level ORFs throughout.

### Gene sets

neORFs were compared to six reference sets. (i) Intergenic, repeat-masked ORFs (N = 1,472,446; ℓ₀.₅ = 22 aa): six-frame, stop-to-stop ORFs (≥10 aa, no ambiguous residue, de-duplicated) extracted from the complement of (annotated genes ∪ RepeatMasker intervals) on the *D. melanogaster* autosome arms (dm6 + UCSC dm6 RepeatMasker track with manual coordinate verification). (ii) GC- and length-matched random ORFs: sequences generated at the intergenic GC content (0.413) and matched to the neORF length distribution, providing a genetic-code null that lacks genuine DNA structure. (iii) Length-matched proteome (N = 7,849): each neORF was matched with a nearest-length protein from the *D. melanogaster* proteome (FlyBase release dmel r6.61; dm6/BDGP6). (iv) Known metal-binding proteins (N = 856; ℓ₀.₅ = 538 aa): UniProt reviewed *D. melanogaster* proteins annotated with the Metal-binding keyword (KW-0479) (FlyBase release dmel r6.61, dm6/BDGP6)(*92*). (v) Drosophila-restricted “young” genes (N = 3042) and (vi) evolutionarily “older” genes (N = 8826). Gene age was assigned by orthology: for each *D. melanogaster* gene, ortholog partners were taken from the NCBI Orthologs resource and each partner mapped to a phylostratum from its NCBI Taxonomy lineage: (1) *melanogaster* group/subgroup, (2) *Drosophilidae* (*Sophophora*/*Drosophila*), (3) *Diptera*, (4) *Insecta*, (5) *Arthropoda* or older, with (0) for genes lacking any ortholog outside *D. melanogaster*. Each gene was assigned the oldest stratum among its partners. “Young” genes are those restricted to *Drosophilidae* or younger (stratum ≤ 2) together with the experimentally defined *de novo* genes of (*93*). “Older” genes have an ortholog at *Insecta* or deeper (stratum ≥ 4). Genes whose oldest ortholog is *Diptera* (stratum 3) fall in neither class. For these annotated sets the longest current FlyBase isoform per gene was used.

### Bis-histidine spacing (order-fold)

For each sequence set, H-(x)ₙ-H configurations (two histidines separated by n intervening residues, n = 1…6) were counted across all sequences. To isolate residue *order* from amino-acid *composition*, the pooled observed count at each n was divided by its expectation under a within-sequence positional null. Each sequence’s own residues were randomly permuted in pace, which preserves the sequence’s length and exact amino-acid composition, randomizing only the positions of histidines. The expectation was taken as the mean over k=3000 such composition-preserving per-sequence scrambles, with the ratio being used to calculate the order-fold value. A difference in pooled order-fold between neORFs and intergenic ORFs was assessed by a two-sample, unpaired bootstrap over sequences (B=3000). In addition, the spacing signal was tested as a per-position 2×2 Fisher’s Exact test, and motif incidence at each spacing was tested by Fisher’s Exact against two composition-aware nulls: (1) random ORFs generated at the intergenic GC content and matched to the neORF length distribution, and (2) a dinucleotide-preserving shuffle of each neORF, which holds dinucleotide content, and thus (CA)n frequency, constant. The identical procedure was applied to C-(x)n-C configurations.

### Motif incidence and density

For H-x-H, C-x-C, and C-x-x-C motifs, incidence (fraction of proteins with ≥1 motif) and density (motifs per residue, i.e. length-normalized) were computed and reported across the random, length-matched-proteome sample, intergenic, neORF, young, old, and metal-binder sets. Because the sets differ ∼25-fold in median length, the incidence and density were reported separately. Incidence differences were tested by 2×2 Fisher’s Exact test (neORF vs each gene class), and density differences were tested by a two-sided Mann-Whitney U test on per-sequence densities. 95% confidence intervals on density were obtained by bootstrap resampling over sequences (B=2000). Per-motif values are presented in Dataset S2.

### Codon usage and the (CA)n mechanism

Histidine is encoded by CAC or CAT; a (CA)n tract read in the His-Thr frame yields CAC·ACA = His-Thr-His, so a (CA)n origin predicts an excess of CAC (over CAT) specifically at motif histidines. Each histidine was assigned to exactly one position (motif vs. non-motif), while histidines that fall in more than one overlapping H-x-H (the central H of an H-x-H-x-H run) were counted once. CAC-vs-CAT usage at motif vs. non-motif histidines was then tested by a single 2×2 Fisher’s exact test (motif/non-motif × CAC/CAT, unique histidine positions), and the same comparison was made against the intergenic-ORF histidine pool. The amino-acid identity at the central (x) position of H-x-H motifs (the (CA)n frame predicts threonine) was tested by a binomial exact test of the observed threonine count against its background frequency in the same set. RepeatMasker was applied before ORF extraction, and short chelation-range (CA)n tracts (n ≈ 7–9) were verified to fall below the RepeatMasker length threshold. (CA)n tracts were thus found to be retained in both the neORF and intergenic sets, so the codon-usage contrast is not a masking artifact.

### Short linear motifs and composition

neORFs and the length-matched proteome were scanned for the Eukaryotic Linear Motif (ELM) classes using the ELM regular-expression catalog (*37*), ranking classes by the difference in per-length density between neORFs and the proteome (Figure S2); H-x-H is not itself an ELM class. Per-class amino-acid composition (histidine, cysteine, aspartate, glutamate, and the full 20-residue profile) was computed across all sequence sets (Figure S1). Per-residue prevalence differences between sets were tested by a binomial test of each set’s residue count against the intergenic residue frequency.

### RNA-seq data and analysis

Expression was quantified from a publicly available *Drosophila* S2R+ RNA-seq series comprising control, zinc-, and manganese-exposed samples (GSE99332; *52*). Reads were aligned to the *D. melanogaster* genome (dm6/BDGP6)(*94*) with STAR (*95*), retaining only uniquely mapped reads, and were counted over neORF and annotated-gene intervals with featureCounts (*96*). Counts were then converted to TPM using ORF length. Two quality-control filters guarded against microsatellite- and repeat-driven mapping artifacts. First, neORFs nominally annotated as intergenic and non-overlapping were re-checked against gene and repeat annotations, and those overlapping annotated features or repeats were excluded. Second, per-base unique-mappability was computed across each locus, and low-mappability loci were removed. These filters yielded 472 uniquely mappable, non-overlapping *de novo* ORFs used for all expression estimates. This number subsequently fell to 466 as 6 ORFs did not return featureCounts values due to near-zero expression.

Differential expression between metal-exposed and control samples was assessed with DESeq2 (*97*) under a genotype-plus-condition design, with independent filtering and Benjamini– Hochberg correction; manganese-versus-control and zinc-versus-control were tested as separate contrasts, and the genome-wide breadth of differential expression was recorded for each. For ZMEG, the S2R+-expressed sequence was reconstructed directly from the mapped reads and compared to the reference assembly. Because dinucleotide microsatellites are prone to alignment-induced indel artifacts, the (CA)n tract length and the open reading frame were determined by manual inspection of the raw reads rather than from the reference alignment. Upstream regulatory content was assessed by scanning 1,200-bp windows around each candidate for MTF-1 metal-response elements (MRE core TGCRCNC, both strands) and canonical core-promoter motifs, calibrated against a null distribution of random intergenic windows and against the metallothionein (MtnA–E) promoters as positive controls.

### Structure and metal-site prediction

The ZMEG open reading frame, and the S100A9 subunit of human calprotectin as a control, were folded with ESMFold (*53*) in the apo state (Figure S4); per-residue confidence is reported as pLDDT. Candidate metal sites and per-voxel zinc-binding probability were predicted with Metal3D (*54*), and the maximum predicted probability p(zinc) and the distance from the highest-probability voxel to the His32–His36 midpoint were recorded. Inter-histidine and metal– nitrogen distances were measured on the apo model with a Zn^2+^ placed for illustration only. Calprotectin was also analyzed identically.

### Idealized-symmetry homodimer construction and scoring

Idealized symmetric C2 homodimers were constructed for ZMEG, DK14275, and ES14265 as follows. A tetrahedral Zn template was defined (Zn at the origin, four coordinating nitrogens at 2.05 Å on tetrahedral vertices) and ESMFold monomers were placed by rigid-body superposition so that one coordinating His pair’s imidazole nitrogens occupied two adjacent vertices, and the second protomer was generated as the C2 image to fill the remaining two. The single residual degree of freedom (rotation of protomer A about the His–His axis) was scanned in 5° increments and the orientation minimizing inter-chain heavy-atom clash (atom pairs < 2.5 Å) retained. Zn was then removed and the apo dimer written for scoring. As ZMEG was the initial test case, its known bis-histidine motif (His32/His36) was used as the specified coordinating pair. The same procedure was then applied to DK14275 and ES14265, for which the coordinating pair was chosen automatically as the closest imidazole-nitrogen pair in the monomer (His54/His58 and His40/His44, respectively).

For O433, the homodimer was predicted directly with Boltz-2 (v2.2.1)(*56*), MSA-free single-sequence, as two identical chains plus one Zn²⁺ (CCD ligand), with five diffusion samples. The sample presenting the most coordinating atoms within 2.6 Å of the Zn was selected. This recovered a symmetric inter-chain His₂Cys₂ site (Cys62 + His89 contributed by each chain). Coordination propensity was scored for these four dimeric structures using Metal3D (*54*), and dimers were ray-traced using PyMOL.

### Comparative synteny and de novo origin trajectory

Orthologous ZMEG loci were recovered in eight outgroup *Drosophila* genomes (*D. simulans*, *D. sechellia*, *D. mauritiana*, *D. yakuba*, *D. erecta*, *D. ananassae*, *D. pseudoobscura*, and *D. virilis*) (NCBI assemblies GCF_016746395.2, GCF_004382195.2, GCF_004382145.1, GCF_016746365.2, GCA_047116545.1, GCF_017639315.1, GCF_009870125.1, and GCF_030788295.1, respectively) by anchored synteny to the flanking single-copy genes Gr93 and glec, followed by local alignment of the intervening region (BLAST, *98*, MAFFT, *99*). For each species, the ORF from the conserved ATG to the first in-frame stop, its length, histidine content, and (CA)n tract length were extracted directly from the assembly; because these genomes are single assemblies and microsatellite length is assembly-sensitive, non-reference tract lengths and their resulting frames are reported as provisional. Character states ((CA)n length, ORF length, and His-motif status) were polarized on the accepted *Drosophila* species topology to order the events shown in Figure 4C, and upstream sequence was aligned and polarized to distinguish ancestral from lineage-derived positions. Within *D. melanogaster*, the (CA)n tract was additionally genotyped across the iso-female founder strains and its length and frame consequence (in-frame deletion versus frameshift) tabulated (Table S1).

### Use of AI and generative AI software

The author used the generative AI large language model Claude Opus 4.8 (Anthropic) to assist with data retrieval, analysis, figure-generation code, and manuscript editing. The author reviewed, fact-checked, and takes full responsibility for all content.

### Data Availability Statement

All analysis and figure-generation code, a single run-all wrapper, and intermediate result tables are provided in the accompanying code release (https://github.com/ulee-sciscripts/denovo_chelation); raw inputs are retrieved by the included fetch_data.sh from Zenodo record 7322757 (neORFs), FlyBase (proteome/annotation), UniProt (metal-binding proteins), and UCSC dm6 (genome, RepeatMasker), and the eight outgroup assemblies from NCBI (accessions in Materials and Methods).

## Supporting information

DataS1

## Acknowledgments

U.L. was supported by a National Science Foundation Postdoctoral Research Fellowship in Biology (NSF PRFB) award no. 2410289.

## Competing Interests

The author declares no competing interest.

## Supplementary Material

**Figure S1.**
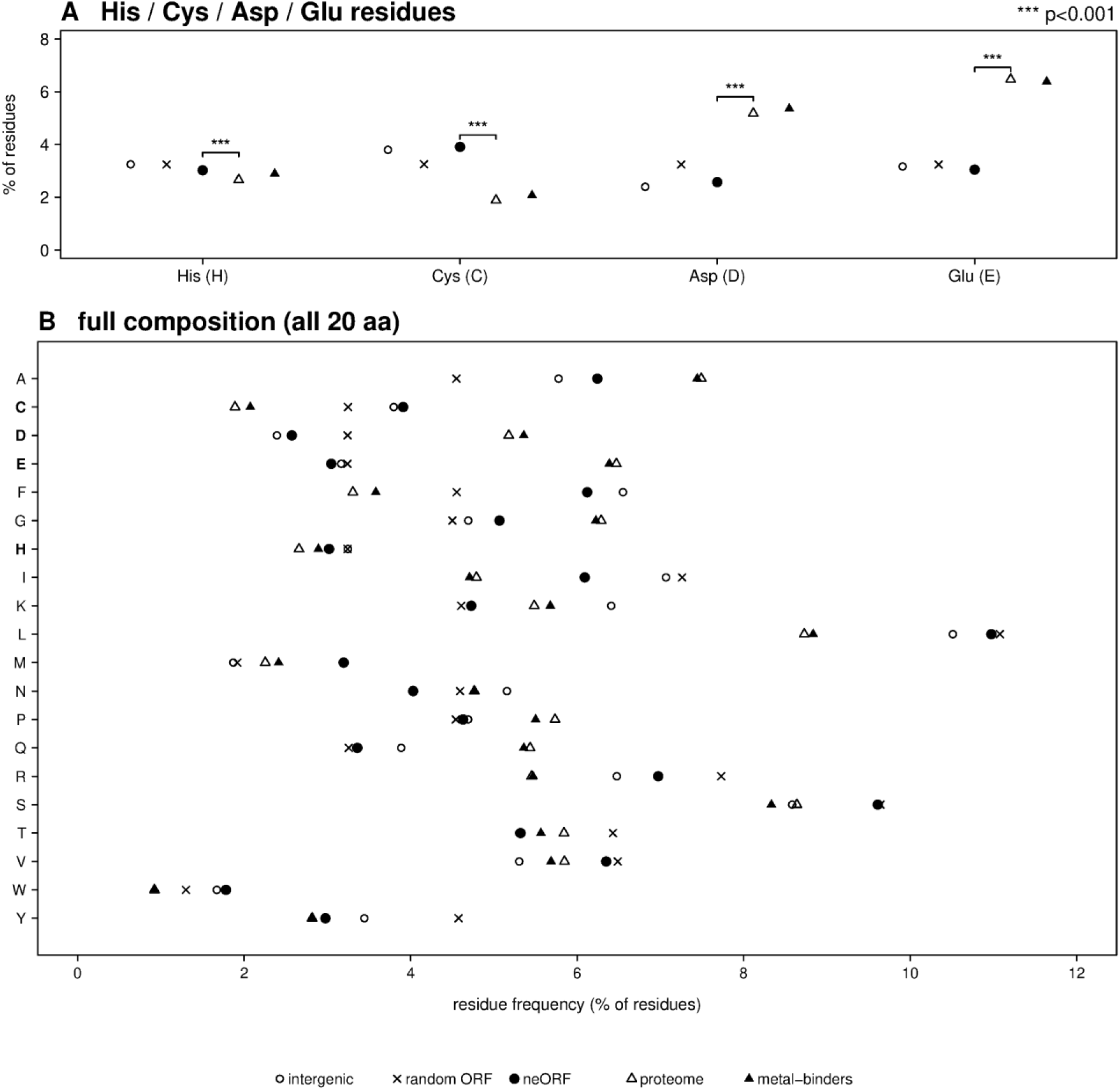
Histidine is not particularly enriched or prevalent in neORFs. Percentage of residues calculated for (A) histidine, cysteine, aspartic acid, glutamic acid, and (B) all amino acids across all gene classes.

**Figure S2.**
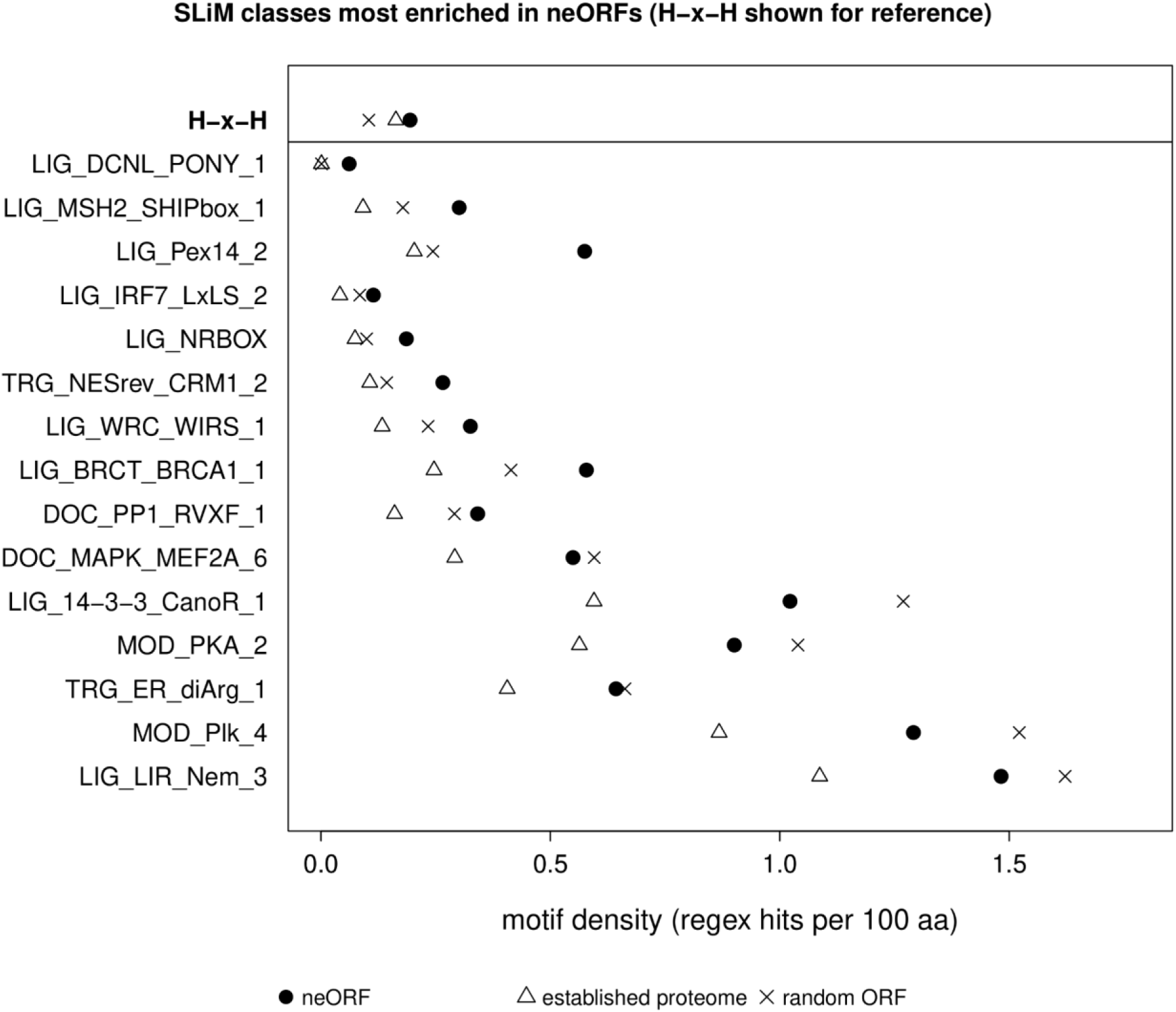
Top 15 most enriched SLiM in neORFs and H-x-H motif prevalence. Motif density for SLiM (ELM catalog) showing largest difference between neORFs and a length-matched proteome sample. Top row shows H-x-H, which was not present in ELM SLiM.

**Figure S3.**
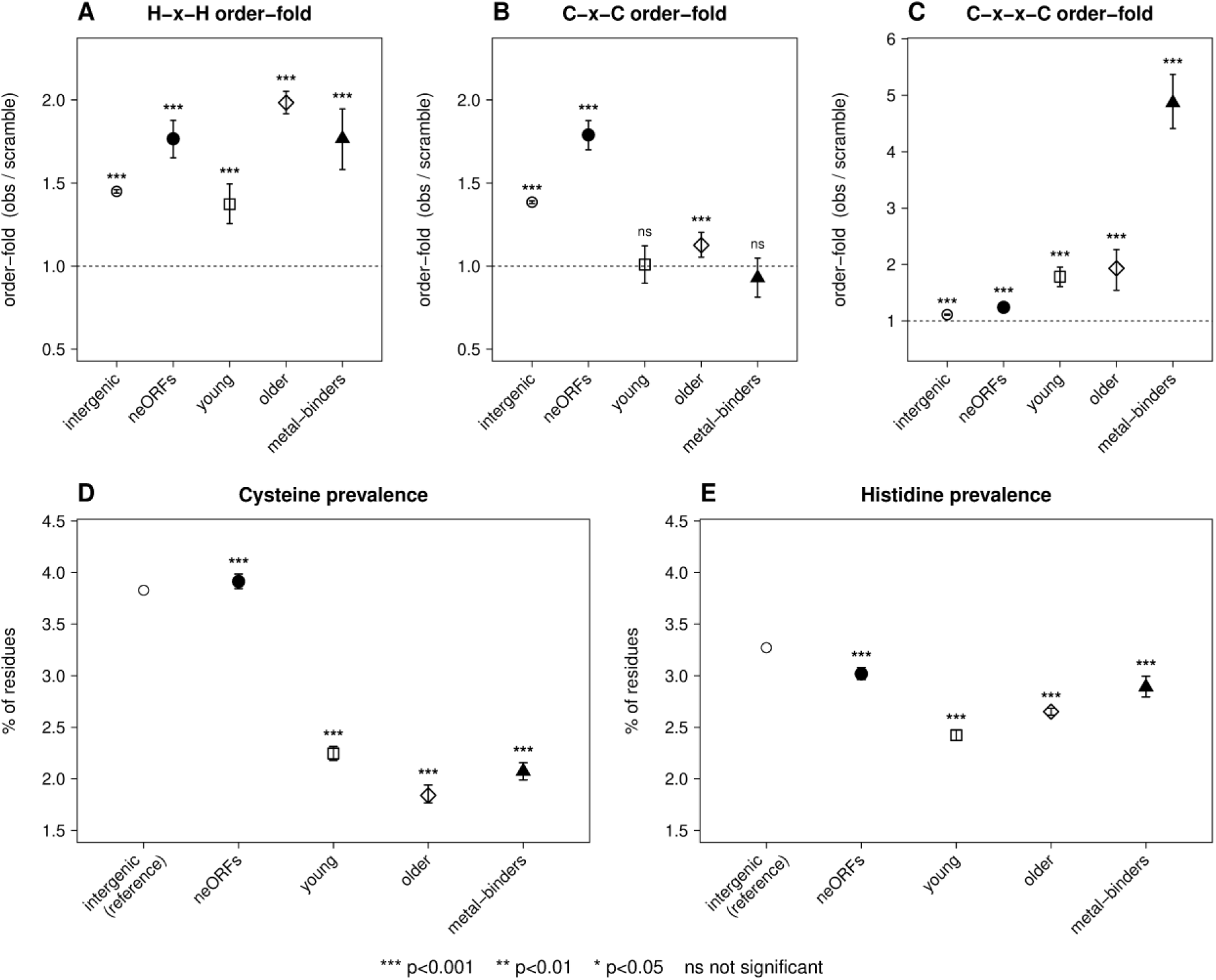
Motif and amino acid prevalence by gene class. Prevalence of (A) H-x-H, (B) C-x-C, and (C) C-x-x-C motifs as well as (D) cysteine and (E) histidine within intergenic, neORFs, young (including duplicate, de novo, and orphan classes), older, and metal-binding gene classes. P-values calculated by (A-C) Fisher’s Exact, or (D-E) binomal test using intergenic genes as a reference.

**Figure S4.**
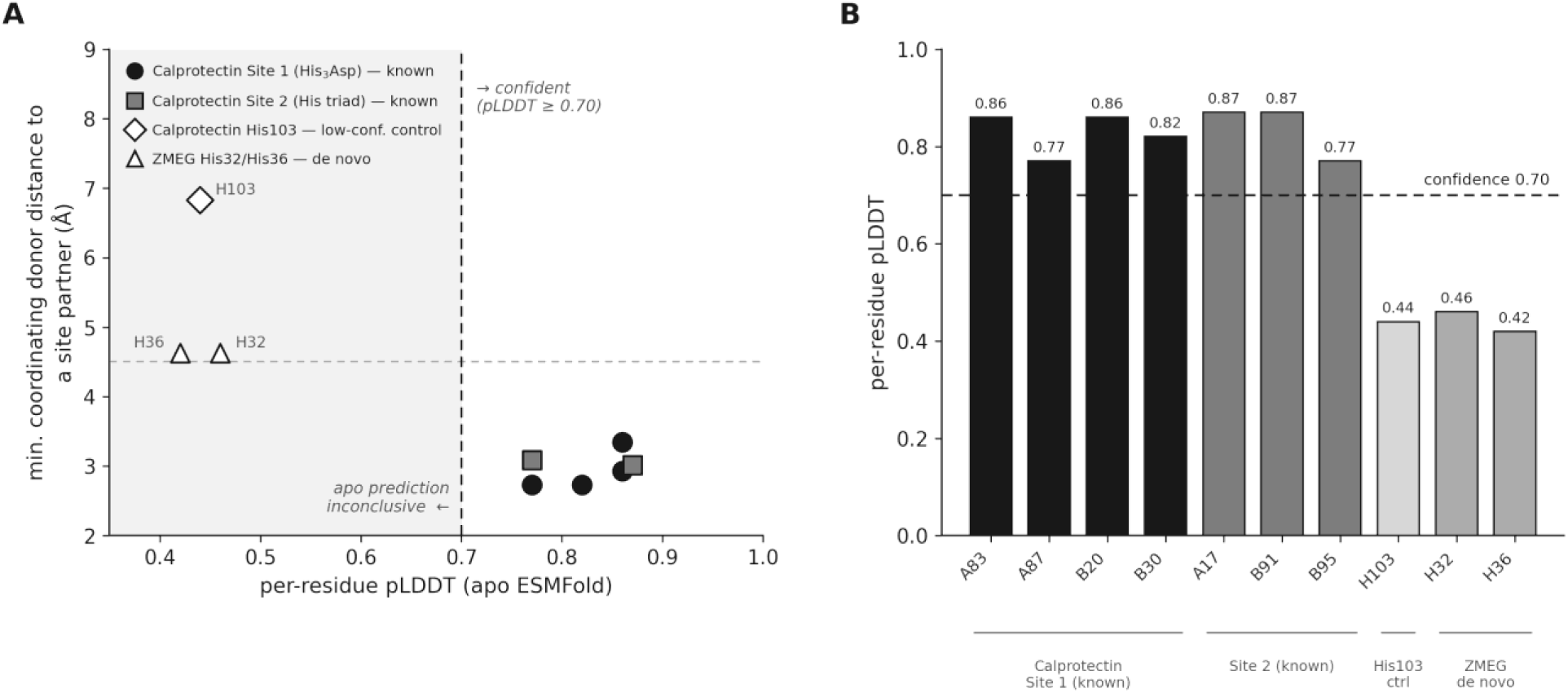
Apo structure prediction does not, on its own, recover calprotectin’s metal site or known coordinating geometry for the S100A9 subunit of calprotectin, whose zinc/manganese site is experimentally established.

*‘Dataset_S1_neORF_expression.csv’*

**Dataset S1.** Per-replicate expression (TPM) of the 466 quantified, uniquely mappable, non-overlapping de novo neORFs in S2R+ cells across control, zinc, and manganese conditions and three genotypes, with H-x-H motif content and dm6 coordinates.

**Table S1.**
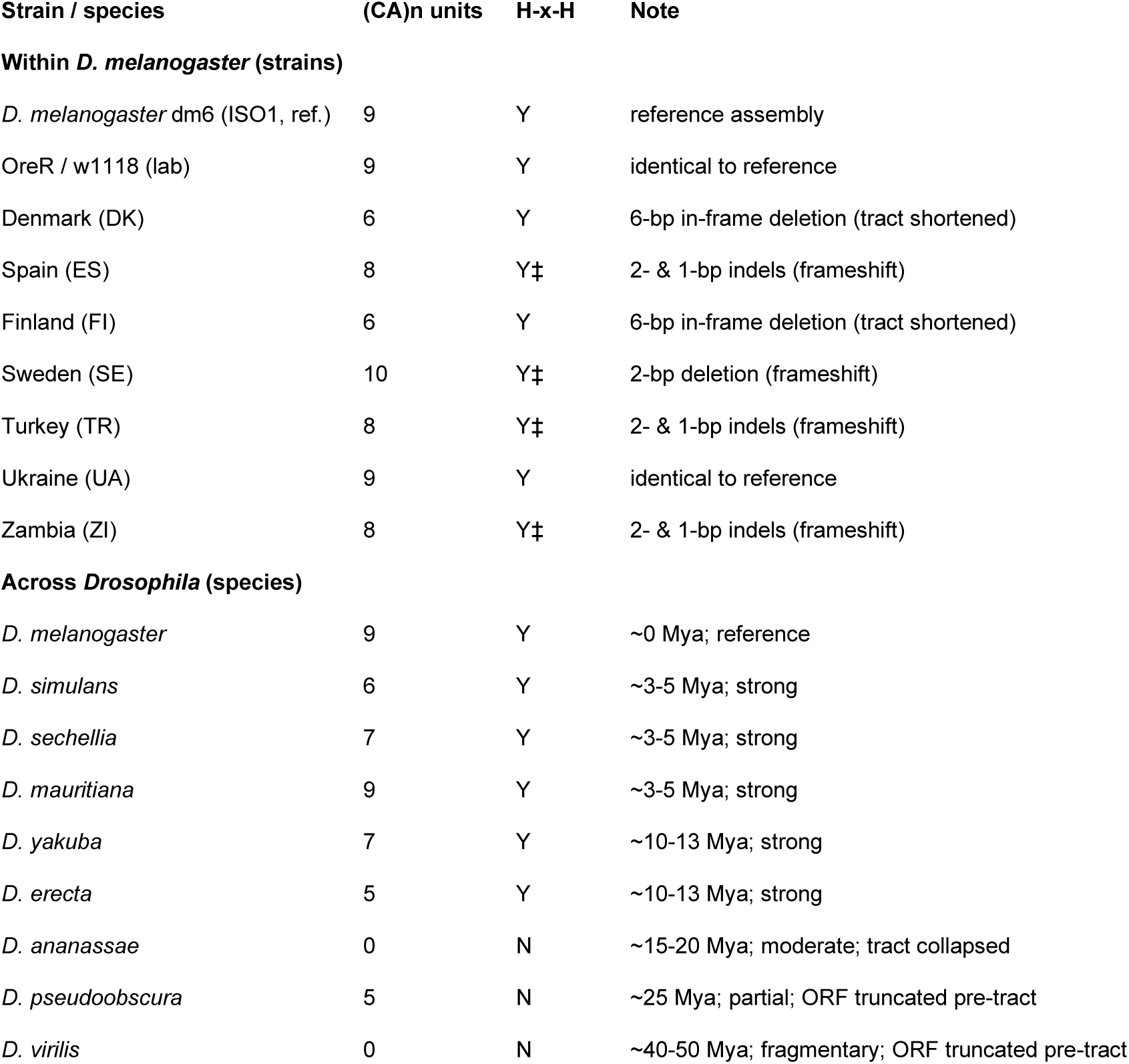
(CA)n tract length and His-motif status at the ZMEG locus within *D. melanogaster* and across *Drosophila*. *‡, His motif present but the (CA)n tract is frameshifted (in-frame His run disrupted)*.

| Strain / species | (CA)n units | H-x-H | Note |
| --- | --- | --- | --- |
| **Within *D. melanogaster* (strains)** |  |  |  |
| *D. melanogaster* dm6 (ISO1, ref.) | 9 | Y | reference assembly |
| OreR / w1118 (lab) | 9 | Y | identical to reference |
| Denmark (DK) | 6 | Y | 6-bp in-frame deletion (tract shortened) |
| Spain (ES) | 8 | Y‡ | 2- & 1-bp indels (frameshift) |
| Finland (FI) | 6 | Y | 6-bp in-frame deletion (tract shortened) |
| Sweden (SE) | 10 | Y‡ | 2-bp deletion (frameshift) |
| Turkey (TR) | 8 | Y‡ | 2- & 1-bp indels (frameshift) |
| Ukraine (UA) | 9 | Y | identical to reference |
| Zambia (ZI) | 8 | Y‡ | 2- & 1-bp indels (frameshift) |
| **Across *Drosophila* (species)** |  |  |  |
| *D. melanogaster* | 9 | Y | ~0 Mya; reference |
| *D. simulans* | 6 | Y | ~3-5 Mya; strong |
| *D. sechellia* | 7 | Y | ~3-5 Mya; strong |
| *D. mauritiana* | 9 | Y | ~3-5 Mya; strong |
| *D. yakuba* | 7 | Y | ~10-13 Mya; strong |
| *D. erecta* | 5 | Y | ~10-13 Mya; strong |
| *D. ananassae* | 0 | N | ~15-20 Mya; moderate; tract collapsed |
| *D. pseudoobscura* | 5 | N | ~25 Mya; partial; ORF truncated pre-tract |
| *D. virilis* | 0 | N | ~40-50 Mya; fragmentary; ORF truncated pre-tract |

*‘Dataset_S2.csv’*

**Dataset S2.** Per-protein H-x-H / C-x-x-C cluster counts and codon usage for neORFs, the *Drosophila melanogaster* proteome, and random ORFs.

## Notes

### Competing Interest Statement

The authors have declared no competing interest.

### Summary of Updates

new population genetics analysis + 2 new sections regarding (TG)n, C-x-C, and O433

