## Supplementary Text for "Naturally arising *de novo* open reading frames as potential zinc chelators in *Drosophila melanogaster*"

### 1 Model

Suppose a recurring cellular demand $D$ for $Zn^{2+}$-buffering capacity is met with the aggregate output of a large pool of $N_{\text{pool}}$ independent, non-linked loci that are transcribed and translated. Assuming that overall enzymatic activity is additive with no deleterious, off-target protein-protein interactions, we may define the genome's aggregate $Zn^{2+}$-buffering capacity $C$ as,

$$C \;=\; \sum_{i=1}^{N_{\text{pool}}} c_i, \tag{1}$$

where $i$ indexes the $N_{\text{pool}}$ contributing loci. We may define fitness as a continuous, once differentiable function whose value depends on the difference between the genome's aggregate capacity $C$ and the demand $D$ resulting from existing environmental conditions such that

$$w \;=\; f(C-D). \tag{2}$$

**Marginal per-locus selection.** Where the genome is well-adapted to local conditions, i.e., the reservoir comfortably meets demand so that $f'(C-D)$ is small (though nonzero), we may consider a single mutation in locus $i$ that alters $c_i$, and thus $C$, by $\Delta c_i$. Such mutations may include, but are not limited to, slippage events, frameshifts, substitutions, or other heritable activating or disabling events at locus $i$. To first order, the change in fitness resulting from such mutation is

$$\Delta w_i = f'(C-D)\,\Delta c_i \;+\; O(\Delta c_i^2). \tag{3}$$

Re-writing the leading term of Eq. (3) as the identity

$$\Delta w_i \;\approx\; \underbrace{f'(C-D)\,C}_{\text{(I)}} \cdot \underbrace{\frac{c_i}{C}}_{\text{(II)}} \cdot \underbrace{\frac{\Delta c_i}{c_i}}_{\text{(III)}}, \tag{4}$$

we may see that the change in fitness is related to (I) demand saturation, (II) functional redundancy, and (III) the intrinsic effect of mutation $i$.

### 2 Per-locus neutrality below the drift barrier

In a genome that is well-adapted to local conditions, and a large number of similarly functioning loci contribute to $C$, (I) and (II) are small. Specifically, following Hartl–Dykhuizen–Dean [1], such a genome operates where fitness is nearly insensitive to further capacity, $f'(C-D) \approx 0$, so any single locus's marginal change in $C$ barely moves fitness. Similarly, the assumption that all loci function similarly does not allow any single locus to carry much more than its fractional contribution $1/N_{\text{pool}}$, where $1/N_{\text{pool}} \to 0$ for large $N_{\text{pool}}$. As a result, the marginal fitness effect of locus $i$ is small in both structural factors ($f'(C-D)\,C$ by saturation and $1/N_{\text{pool}}$ by redundancy) even when $|\Delta c_i/c_i|$ is of order-of-magnitude 1, as

$$|\Delta w_i| \;\approx\; f'(C-D)\,C \cdot \frac{1}{N_{\text{pool}}} \cdot \left|\frac{\Delta c_i}{c_i}\right|. \tag{5}$$

This relationship demonstrates how a single mutation in locus $i$ is effectively neutral once its coefficient falls below the nearly-neutral drift barrier [2, 3, 4]

$$|\Delta w_i| \;\lesssim\; \frac{1}{2N_e}. \tag{6}$$

Thus, for *D. melanogaster*, with $N_e \approx 1 — 2.5 \times 10^6$ [6, 7], the barrier is

$$\frac{1}{2N_e} \;\approx\; 2\times 10^{-7} — 5\times 10^{-7}. \tag{7}$$

Assuming a pool of $N_{\text{pool}} \approx 7{,}000$ *de novo* loci [5], the product $f'(C-D)\,C \cdot \left|\Delta c_i/c_i\right|$ would need to exceed $\approx 1.4\times 10^{-3} — 3.5\times 10^{-3}$ to clear this barrier. Consequently, even if the entire buffering function carried a modest marginal fitness weight $f'(C-D)\,C$ of order 0.1%, a single locus would need to change its own output $\left|\Delta c_i/c_i\right|$ by a full 140% — 350% to be readily selectable in *D. melanogaster*. Since abolishing an entire locus is only a 100% change, no disabling mutation at a single locus clears the barrier.
